# High-precision neurofeedback-guided meditation training optimises real-world self-guided meditation practice for well-being

**DOI:** 10.1101/2024.10.25.618656

**Authors:** Saampras Ganesan, Nicholas T. Van Dam, Sunjeev K. Kamboj, Aki Tsuchiyagaito, Matthew D. Sacchet, Masaya Misaki, Bradford A. Moffat, Valentina Lorenzetti, Andrew Zalesky

**Affiliations:** Department of Biomedical Engineering, The University of Melbourne, Australia; Department of Psychiatry, The University of Melbourne, Australia; Contemplative Studies Centre, Melbourne School of Psychological Sciences, The University of Melbourne, Australia; Research Department of Clinical, Educational and Health Psychology, University College London, United Kingdom; Neuroscience of Addiction and Mental Health Program, Healthy Brain and Mind Research Centre, School of Behavioral and Health Sciences, Faculty of Health, Australian Catholic University, Australia; Melbourne Brain Centre Imaging Unit, Department of Radiology, The University of Melbourne, Australia; Laureate Institute for Brain Research, Tulsa, OK, USA; Meditation Research Program, Department of Psychiatry, Massachusetts General Hospital, Harvard Medical School, Boston, Massachusetts, USA; The University of Tulsa, Oxley College of Health & Natural Sciences, Tulsa, OK, USA

**Keywords:** meditation, neurofeedback, 7 Tesla functional MRI, emotional well-being, awareness, posterior cingulate cortex

## Abstract

Meditation can benefit well-being and mental health, but novices often struggle to effectively recognize and disengage from mental processes during meditation due to limited awareness, potentially diminishing meditation’s benefits. We investigated whether personalised high-precision neurofeedback (NF) can improve disengagement from mental activity during meditation and enhance meditation’s outcomes. In a single-blind, controlled, longitudinal paradigm, 40 novice meditators underwent two consecutive days of meditation training with intermittent visual feedback from either their own (N=20) or a matched participant’s (N=20; control group) posterior cingulate cortex (PCC) activity measured using 7 Tesla functional magnetic resonance imaging. During training, the experimental group showed stronger functional decoupling of PCC from dorsolateral prefrontal cortex, indicating better control over disengagement from mental processes during meditation. This led to greater improvements in emotional well-being and mindful awareness of mental processes during a week of real-world self-guided meditation. We provide compelling evidence supporting the utility of high-precision NF-guided meditation training to optimise real-world meditation practice for well-being.

## INTRODUCTION

Meditation is a widespread contemplative practice that involves training attention and cultivating active and receptive awareness of thoughts, emotions, sensations, and mind-body perceptions *(1)*. Meditation can enhance emotion regulation, weaken maladaptive psychological patterns, and promote overall well-being, offering transdiagnostic benefits in alleviating pervasive mental health and mood disorders *(2–4)*. A widely practised and studied meditation technique, i.e., focused attention meditation, typically instructs one to focus on bodily breathing sensations, recognize and disengage from mental distractions when they arise, and refocus on breathing *(5)*. Developing attentional skills through this technique is crucial for progressing to more advanced meditation practices *(6–8)*. However, novices are often deeply entrenched in self-referential mental processing *(84)* with limited meta-cognitive awareness and attention regulation capacity, which can contribute to limited recognition of and disengagement from mental distractions during meditation *(9*, *10)*. These factors can diminish the quality and psychological benefits of meditation practice, discouraging continued practice *(11–14)*. Currently, hundreds of hours of self-guided practice may be necessary to achieve significant relief from psychological distress *(3)*. Furthermore, the subjective nature of meditation and its instructions can complicate the monitoring of meditation practice *(15*, *16)*.

Integrating objective neurotechnology like neurofeedback (NF) with meditation training can potentially contribute to addressing some of the challenges experienced in meditation by novice meditators. NF is an approach that delivers real-time feedback on personalised brain activity that can assist trainees in learning to self-regulate desired mental states, behaviours, or pathologies *(17)* (**Figure 1**).

**Figure 1:**
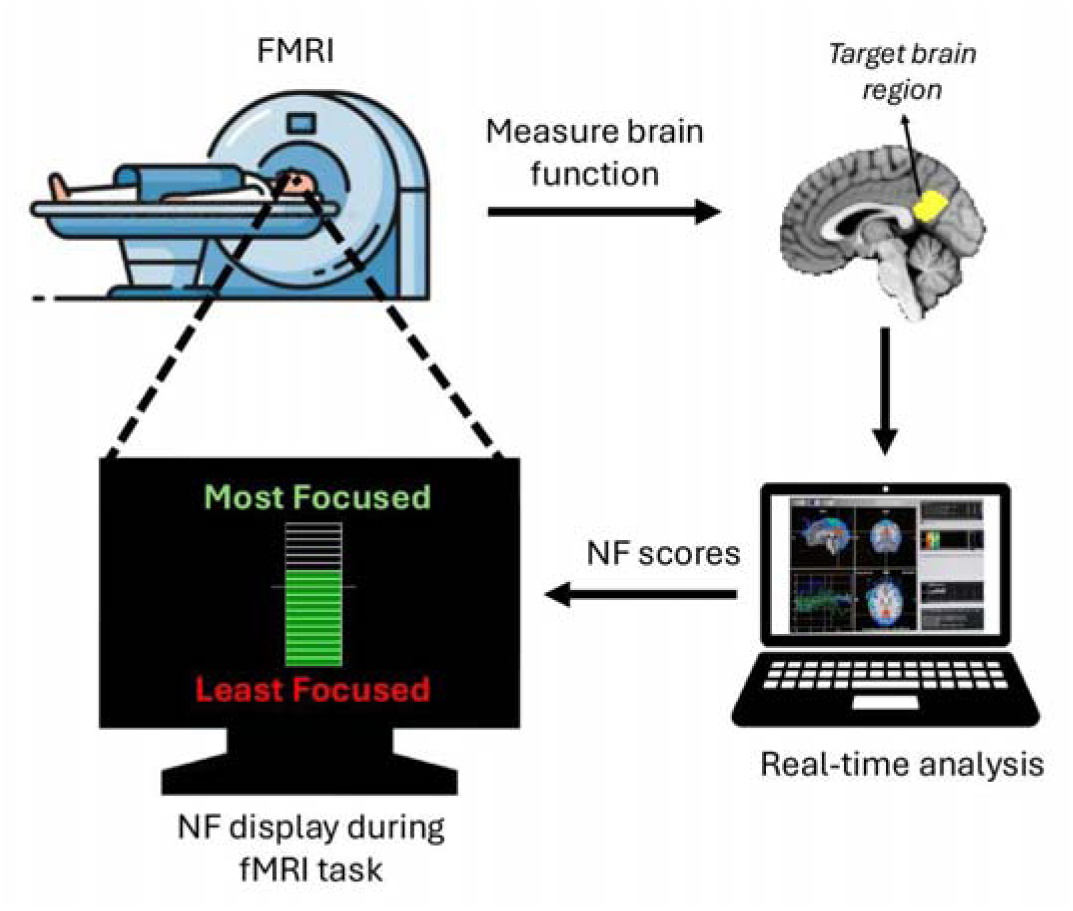
Schematic of real-time functional MRI (fMRI) neurofeedback (NF). The participant’s brain function is measured and analysed in real-time using fMRI, generating NF scores that are displayed to the participant (via a thermometer with incremental bars) to facilitate self-regulation of the target brain function in a closed-loop. In the current study, the posterior cingulate cortex (PCC), shown in yellow, was used as the target brain region.

Emerging evidence suggests that NF training likely leverages reinforcement learning *(18)*, with capacity to enduringly impact behaviour and brain function in healthy individuals, while also aiding in the normalisation of brain function in various pathologies *(17*, *19*, *20)*. While NF using electroencephalography (EEG) is more common and cost-effective, NF using functional magnetic resonance imaging (fMRI) offers comparatively superior neuroanatomical resolution, with capacity for more significant positive effects on neural, clinical and behavioural measures after shorter training durations *(17*, *20–23)*. For example, a single fMRI-NF session significantly improved attention in healthy participants compared to control groups receiving sham NF, no NF, and non-neural NF *(24)*. Similarly, a randomised controlled trial (RCT) showed that two fMRI-NF sessions helped patients with depression upregulate amygdala activity, significantly reducing symptoms compared to a sham-NF group *(25)*. The advent of ultra-high field 7 Tesla MRI, offering superior neuroanatomical precision, statistical power, and signal quality compared to 3 Tesla MRI, holds promise for further enhancing the neuroanatomical precision achievable with fMRI-NF *(26*, *27)*.

The posterior cingulate cortex (PCC), within the default-mode network (DMN), has been widely implicated in mental activity and self-referential processing *(28)*, and is often found to reduce its activation (i.e., deactivate) during focused attention meditation relative to a control condition, as indicated by the fMRI literature on focused attention meditation *(6)* as well as expert meditator reports *(29)*. While fMRI-NF with meditation can modulate meditation-related brain targets including the DMN *(29–34)*, its effects on self-guided meditation practice and meditation’s psychological benefits remain unclear.

To address this critical gap, we introduce a controlled, single-blind, longitudinal, proof-of-concept paradigm to assess the impact of multi-day fMRI-NF meditation training on mindful awareness during subsequent real-world meditation practice (i.e., self-guided silent meditation with eyes closed in a desired posture) among healthy novices. We also tested the effect of this post-NF real-world practice on breath counting skill (an objective proxy for mindfulness) and emotional distress - a measure of negative emotional states related to depression, anxiety and stress - in the healthy novices.

## RESULTS

We leveraged 7 Tesla fMRI and employed a comprehensive study design (**Figure 2**) featuring a yoked-sham NF control group (i.e., group receiving feedback derived from matched participants in the experimental group meditating with veridical NF). The PCC was chosen as the NF brain target. The study (**Figure 2A**) included two sessions of 7 Tesla fMRI NF-guided meditation training over two consecutive days. The NF training was followed by mobile ecological momentary assessments (mEMA) to evaluate changes in mindful awareness during a week of real-world practice at home. A 1-week follow-up assessed emotional distress during the practice week, and breath counting skill after the practice week. We hypothesised that participants who completed NF training, compared to the control group, would (i) learn to engage the targeted brain region more effectively, (ii) demonstrate greater improvements in mindful awareness during self-guided meditation practice over one week, (iii) show more significant reductions in emotional distress during the meditation practice week, and (iv) experience greater enhancements in breath counting skill at the end of the practice week.

**Figure 2:**
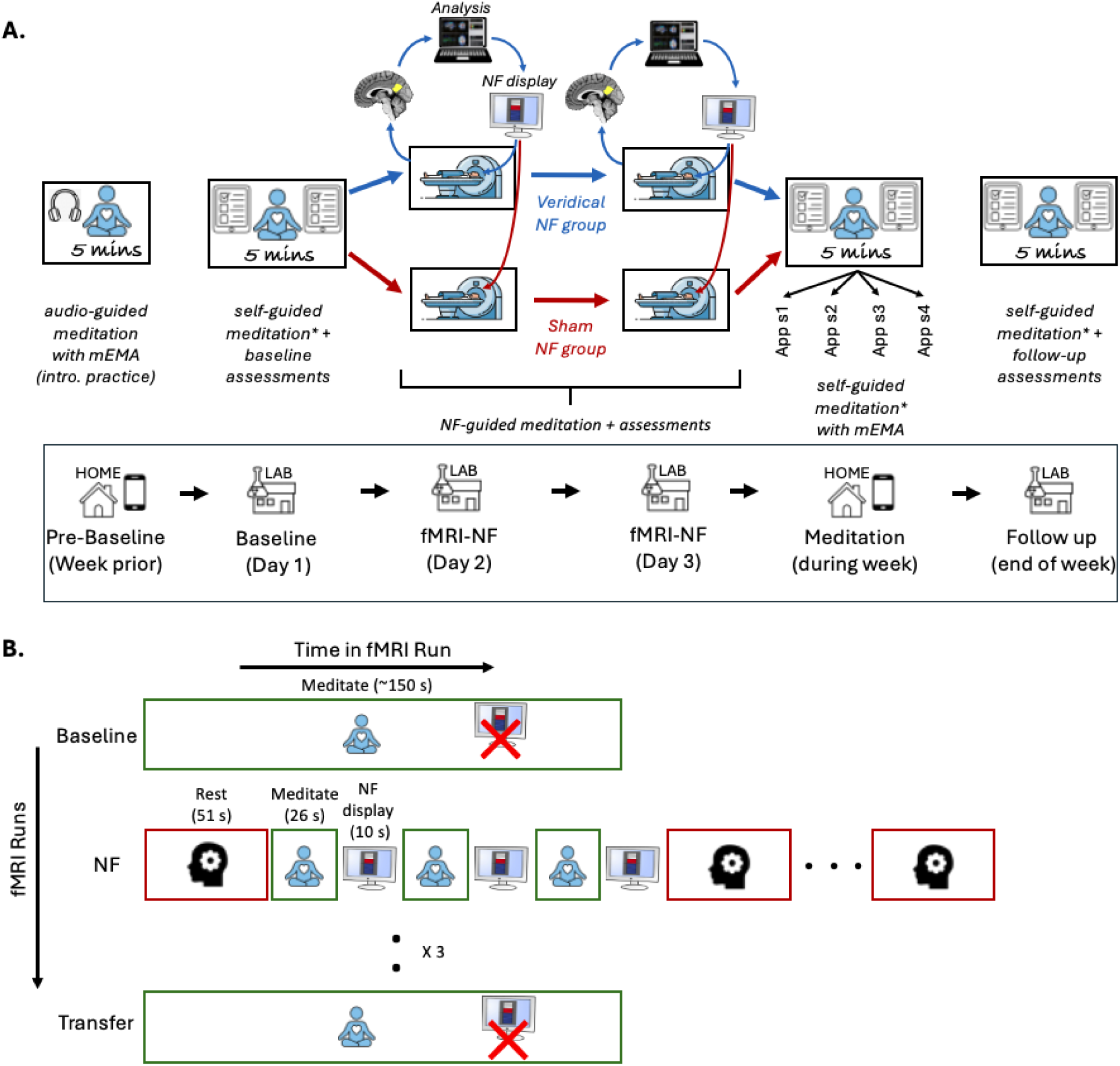
Schematic of the experimental designs used in this study. **A)** The study included a pre-baseline period with introductory audio-guided meditation practice at home using the mEMA app, a baseline session including the first 5-minute self-guided meditation on Day 1, fMRI-neurofeedback (NF) training on Days 2 and 3, a week of self-guided meditation (four 5-minute sessions) at home using the mEMA app, and a follow-up session at the end of the week, which included the final 5-minute self-guided meditation session. A thermometer with incremental bars was used to display real-time visual feedback during the NF sessions. **B)** The fMRI design diagram outlines the NF training procedure, detailing fMRI runs (y-axis) and the conditions/tasks within each run (x-axis). Participants completed a baseline meditation run without NF, three NF runs involving rest (3 blocks) and meditation (6 blocks) with intermittent NF display, and a transfer meditation run without NF to evaluate transfer learning. In each NF run, there were three pairs of meditation and NF blocks between each pair of rest blocks. The same procedure was followed on both NF sessions (Days 2 and 3). mEMA - mobile Ecological Momentary Assessment; NF - Neurofeedback; sx - session x; * Immediately after each 5-minute meditation session, participants completed the self-report State Mindfulness Scale (SMS) which assessed mindful awareness during the session, with pre-session SMS scores serving as a control for baseline mindful awareness.

### Sample and experimental design

40 eligible adults with beginner-level meditation experience and no psychiatric or neurological diagnoses completed the study *(Supplementary Section 1*; **Table 1** for sample characteristics; *Figure S1)*. Participants were assigned to either the experimental group (N=20), which meditated with veridical PCC NF, or a control group (N=20), which meditated with yoked-sham NF based on PCC activity from an experimental participant with similar meditation experience. The participants were blinded to the control group’s existence.

**Table 1:**
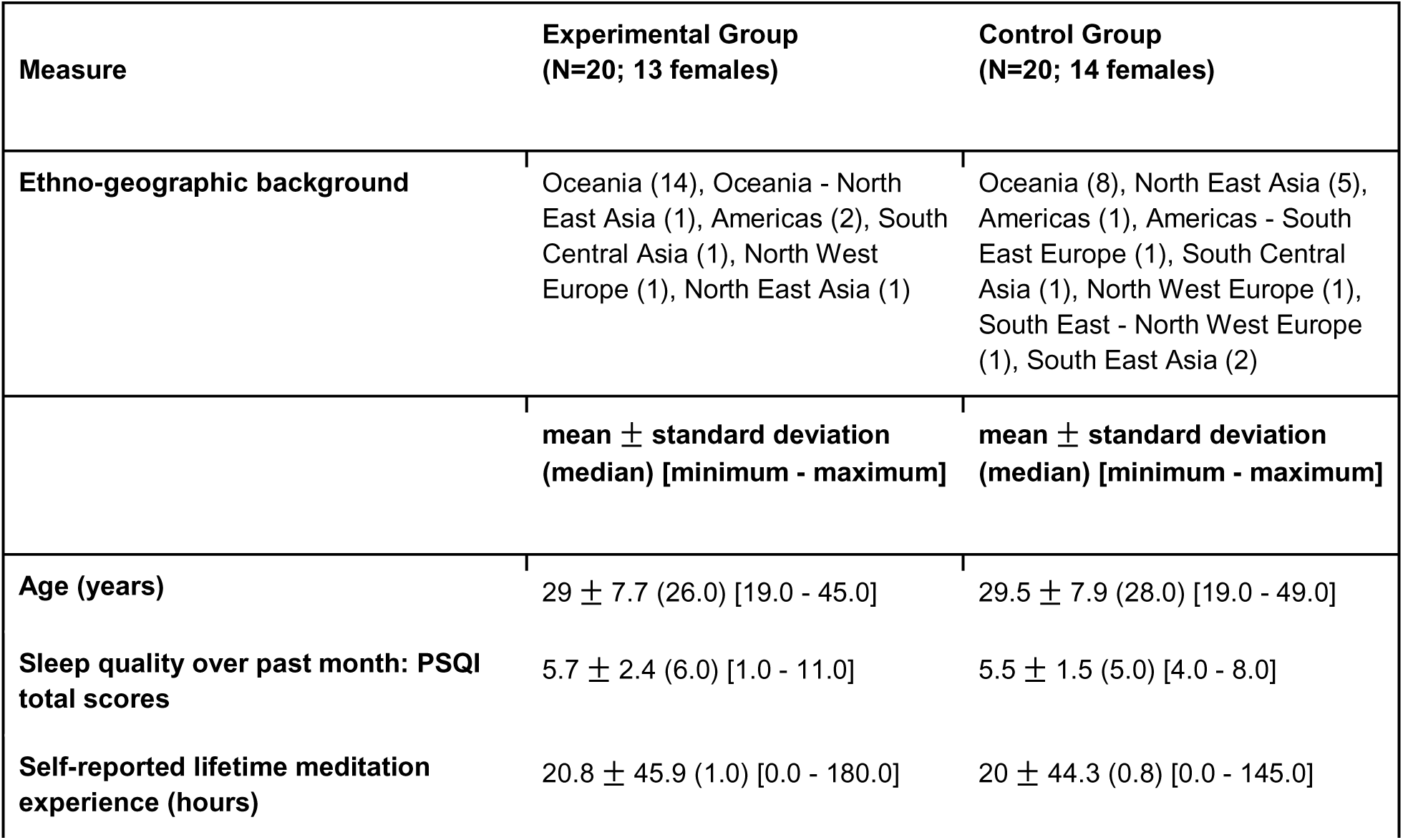
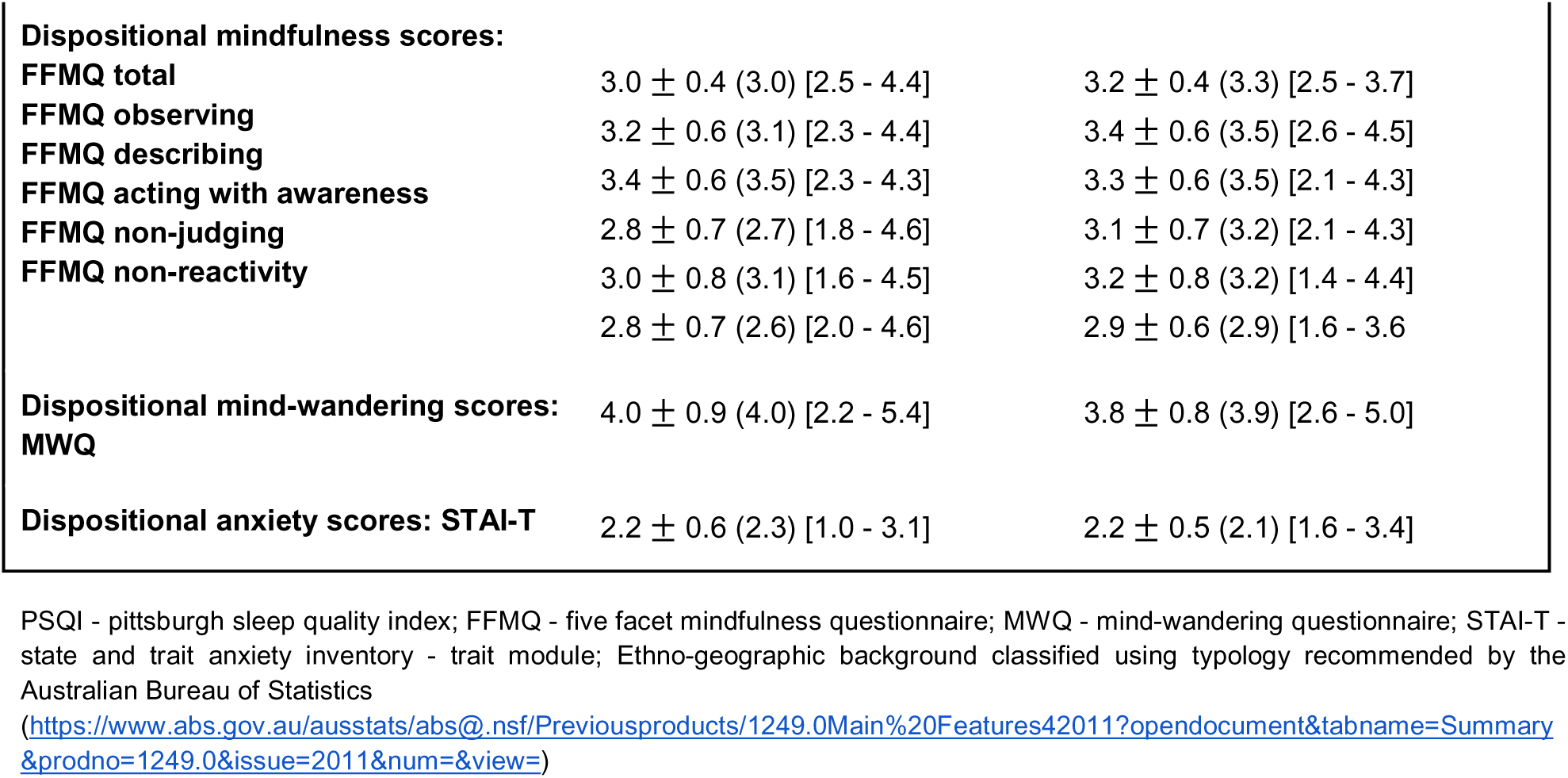
Key sample and baseline characteristics by group. No significant differences were observed between groups.

The groups were equivalent at baseline (**Table 1)**. All participants completed six real-world self-guided 5-minute meditation sessions over one week (one at baseline, four at home post-NF, and one at follow-up). Mindful awareness during each self-guided meditation session was assessed with the State Mindfulness Scale (SMS) *(35)*, and emotional distress over the past week was measured using the self-report Depression, Anxiety, Stress Scale (DASS-21) *(36)* at baseline and follow-up. Stanford Sleepiness Scale (SSS) *(37)* was administered alongside each SMS measurement and after each fMRI run to capture momentary arousal. Additionally, breath counting accuracy during 15-minutes of meditation at baseline and follow-up was evaluated using the Breath Counting Task (BCT) *(38)*.

Each fMRI-NF session included three NF-meditation runs with six 26-second meditation blocks, sandwiched between pre-NF baseline and post-NF transfer meditation runs without feedback (**Figure 2B**). The average percent signal change (PSC) in a dynamic voxel set within the PCC was computed every TR (800ms) using Turbo BrainVoyager software *(Methods)*, relative to a baseline of rest. NF scores derived from PCC PSC averaged within a meditation block were visually displayed *(Figure S2)* to participants after each meditation block, with higher scores indicating more negative PSCs (i.e., greater PCC deactivation).

### Verification of blinding and online processing

The sham PCC signals provided to the control group were not significantly correlated (p>0.05) with their actual PCC signals in either NF session, regardless of whether online or offline fMRI denoising was applied. Furthermore, self-reports on the correspondence between NF scores and meditation, on the utility of NF for learning meditation, and on in-scanner meditation performance were not significantly different between groups in either NF session (≥ 3/5 median rating per group; p>0.05) (**Figure S3A-S3C**), supporting the effectiveness of the blinding. Participants from both groups reported engaging in focused attention meditation anchored in breathing sensations during meditation conditions, and unconstrained thought during rest conditions.

In both NF sessions, PCC deactivation estimated online during NF was significantly correlated with PCC deactivation calculated offline after more stringent preprocessing and denoising (rho = 0.38; p < 0.05; **Figure S3D**). Furthermore, the signal from the control region, which was used as an online proxy for physiological artefacts during NF-guided meditation, did not significantly differ between groups in either session during meditation vs. rest (p > 0.05; **Figure S4**).

### Behavioural outcomes of NF training

#### Mindful awareness during one-week real-world meditation practice

We examined mindful awareness during the week-long post-NF real-world meditation practice using the self-report SMS subscales, which assess awareness of mental activity (SMS-Mind) and bodily sensations (SMS-Body). Specifically, we analysed group differences in the trajectory of change (i.e., slope) in SMS scores linked to mindful awareness during meditation practice comprising six meditation timepoints over one week (1 at baseline, 4 at home, and 1 at follow-up), while controlling for arousal, age, and sex *(Figure S6)*. As the week progressed, the experimental group showed greater increases in mindful awareness of mental activity during meditation practice (more positive SMS-Mind slopes), compared to the control group (Cohen’s *d*=0.38; FDR-adjusted p=0.041; **Figure 3A**), despite equal practice durations. No significant group difference was found for the SMS-Body subscale (**Figure 3B**). Mindful awareness during the baseline 5-minute meditation session did not significantly differ between groups (p>0.05).

**Figure 3:**
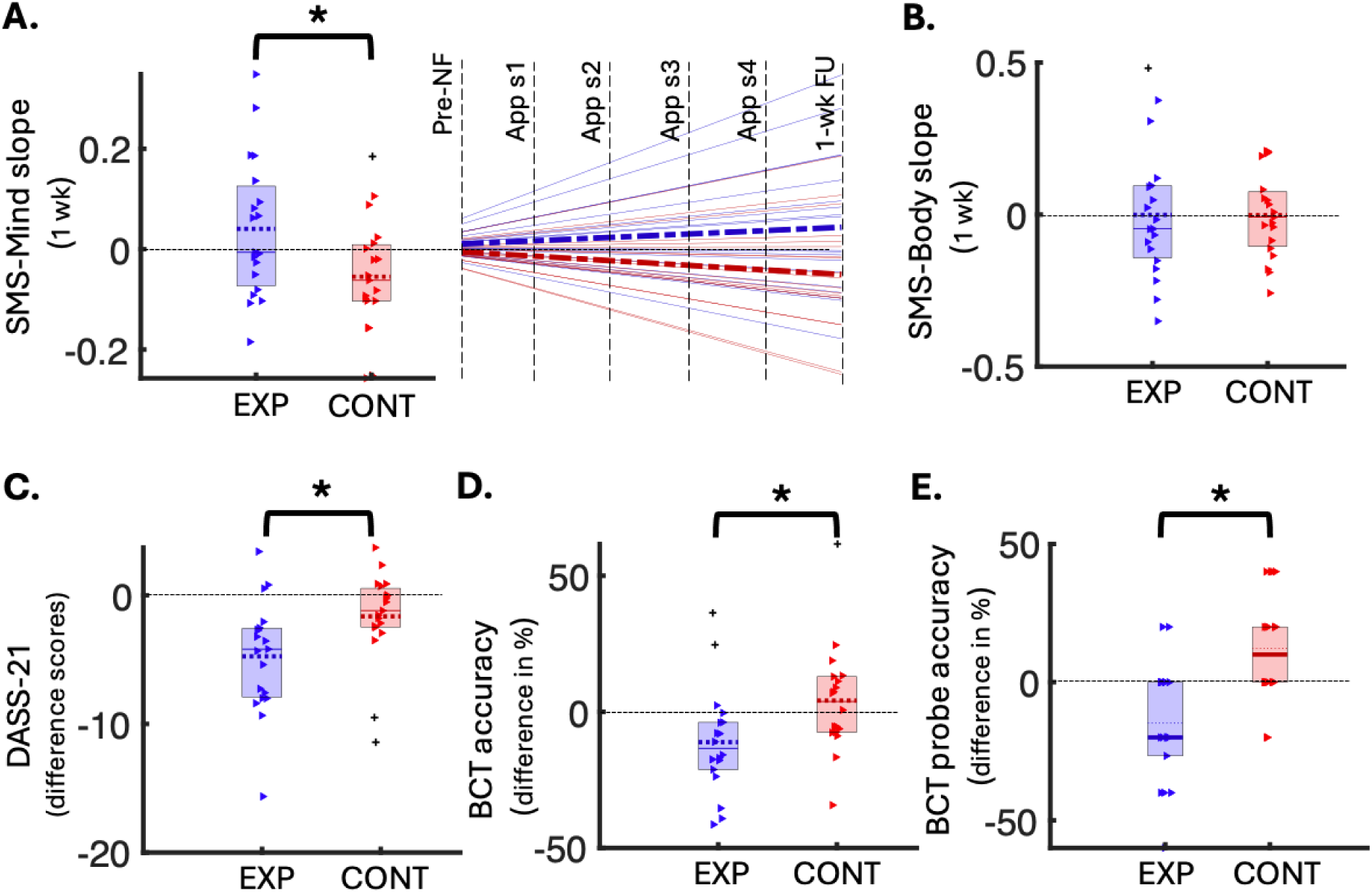
Behavioural outcomes of the NF-guided meditation training. **A)** Left: Box plot of group differences in slopes of change (y-axis) in SMS-Mind scores over 1-week meditation practice (5-minute sessions), significant after FDR correction (FDR-p = 0.041, uncorrected p = 0.027, Cohen’s d = 0.38, N(exp) = 19, N(cont) = 19). Right: Same data represented using a line graph, with the pre-NF data normalised to 0, group mean slopes indicated as bold dashed lines and individual slopes shown as faded solid lines (experimental in blue, control in red). **B)** Box plot of group differences in slopes of change (y-axis) in SMS-Body scores over 1-week meditation practice (5-minute sessions), not significant (N(exp) = 19, N(cont) = 19). **C)** Box plot showing group differences in DASS-total difference scores (y-axis; follow-up minus baseline), significant after FDR correction (FDR-p = 0.041, uncorrected p = 0.025, Cohen’s d = 0.40, N(exp) = 19, N(cont) = 19). **D)** Box plot of group differences in BCT accuracy changes (y-axis; follow-up minus baseline), significant after FDR correction (FDR-p = 0.041, uncorrected p = 0.033, Cohen’s d = 0.39, N(exp) = 18, N(cont) = 18). **E)** Box plot of group differences in BCT probe accuracy changes (y-axis; follow-up minus baseline), significant after FDR correction (FDR-p = 0.005, uncorrected p = 0.001, Cohen’s d = 0.56, N(exp) = 18, N(cont) = 18). In all box plots, coloured triangles represent individual data points (red for control, blue for experimental), with outliers shown as pluses. Means are dotted lines and medians are solid lines. EXP - Experimental group; CONT - Control group; DASS - Depression, Anxiety & Stress Scale; SMS - State Mindfulness Scale; App sx - App meditation session x; wk - week; BCT - Breath Counting Task; NF - Neurofeedback; FU - Follow up; * FDR-significant p<0.05

#### Emotional distress during one-week real-world meditation practice

We investigated the change in emotional distress from the week before baseline to the end of post-NF real-world meditation practice. Specifically, we compared the change in DASS-21 total scores (i.e., follow-up minus baseline) from baseline to one-week follow-up between the veridical and sham NF-guided meditation training groups, controlling for age and sex. The experimental group, compared to the control group, exhibited a significantly greater reduction in emotional distress – indicated by a more negative difference score – from baseline to follow up (Cohen’s *d*=0.40, FDR adjusted p=0.041; **Figure 3C**). The DASS-21 total scores did not differ significantly between groups at baseline (p>0.05).

#### Breath counting after one-week real-world meditation practice

We assessed the change in breath counting accuracy during meditation from baseline to the end of the week involving post-NF real-world meditation practice. Specifically, we assessed group differences in BCT performance by comparing changes in task and probe accuracies from baseline to one-week follow-up (i.e., follow-up minus baseline), controlling for age and sex. Contrary to expectations, the experimental group showed a significant decline (i.e., more negative difference scores) in both BCT task performance indices from baseline to follow up, compared to the control group (task-accuracy FDR-adjusted p=0.041; probe-accuracy FDR-adjusted p=0.005; **Figures 3D and 3E**). A 5% FDR correction was used to adjust for multiple comparisons across all behavioural analyses *(Table S2)*.

The task and probe accuracies were not significantly different between groups at baseline (p>0.05). The breath counting was physiologically valid, as the mean breath rate measured by the respiration belt showed a strong correlation with the mean counting rate (rho=0.98, p<0.0001). Post-hoc exploratory analyses revealed a significant correlation between the increase in mindful awareness during the week-long meditation practice (SMS-Mind slopes) and decrease in BCT task accuracy from baseline to the end of the week-long practice (rho(33)=-0.35, p=0.044; **Figure S5**). Furthermore, at baseline, the BCT task accuracy was not significantly correlated (p>0.05) with dispositional mindfulness (FFMQ and subscales).

### Changes associated with the NF brain target

#### Deactivation during NF-guided meditation

We investigated the effect of NF-guided meditation on deactivation in the NF target region (PCC) (**Figure 4A**). Specifically, we compared mean PCC deactivation during meditation relative to rest between the two groups for each NF session. This offline analysis included stringent control for physiological and head motion artefacts, age, sex and sleepiness. No significant differences in PCC deactivation were observed between the groups in either session (p > 0.05; **Figure 4B**).

**Figure 4:**
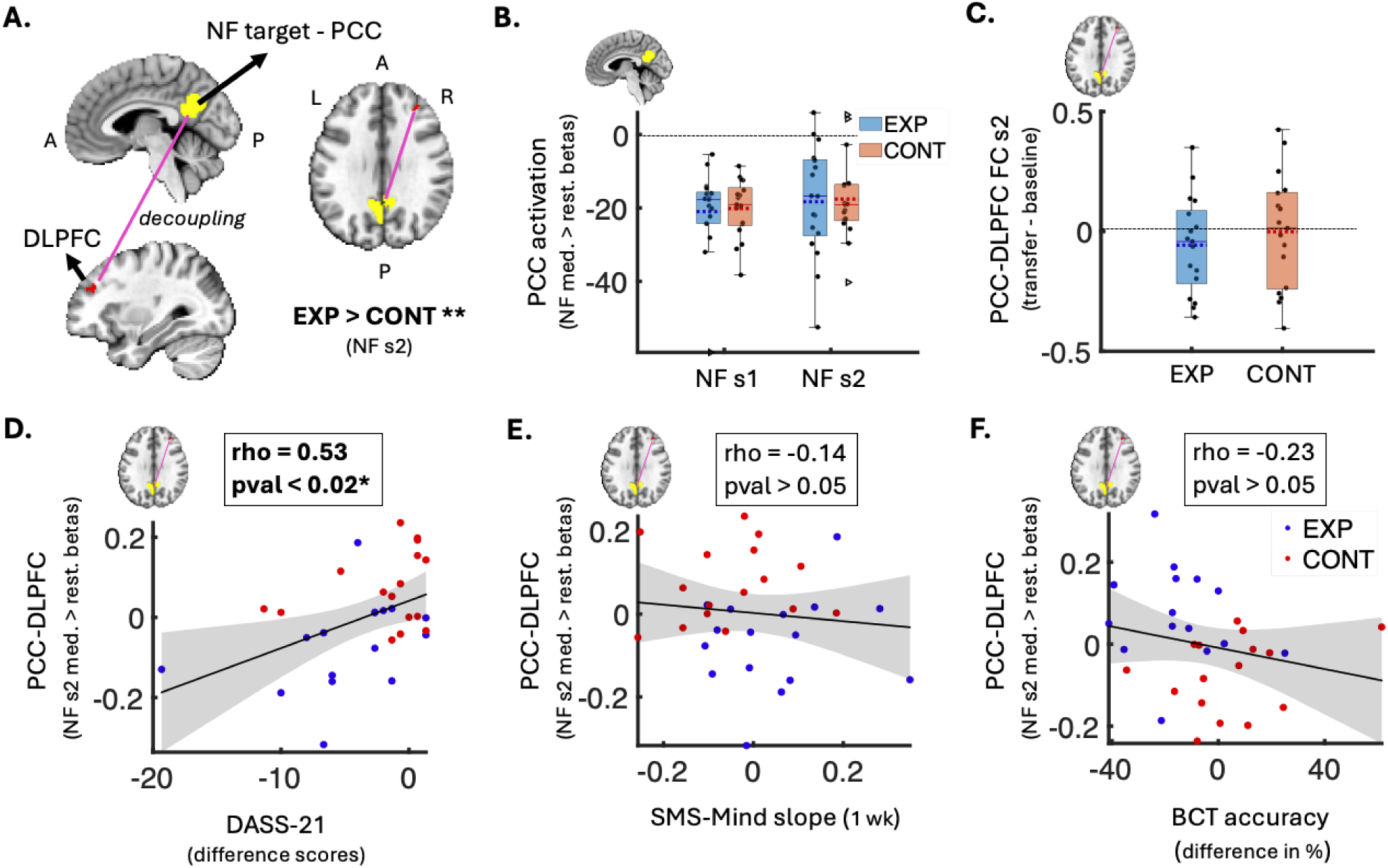
Neural changes and brain-behavior associations related to the NF-guided meditation training. **A)** Brain volume slices showing the decoupling (pink line) between the NF target (PCC in yellow) and right DLPFC (red; gPPI cluster of 26 voxels) that was significantly stronger in the experimental group vs. control group during NF-guided meditation vs. rest (session 2 FWE-p < 0.05, Cohen’s d = 0.59, N(exp) = 17, N(cont) = 17). **B)** Box plot of group difference (not significant) in mean PCC activation betas during NF-guided meditation vs. rest in each NF session (N(exp) = 17, N(cont) = 17). Y-axis represents mean activation values (betas). **C)** Box plot of group difference (not significant) in the change in PCC-DLPFC decoupling (transfer minus baseline) during non-NF meditation in session 2 (N(exp) = 17, N(cont) = 17). Y-axis represents the difference in FC correlation values. Box plots B-C show individual data points as black dots, outliers as triangles, data range as whiskers, means as dotted lines, and medians as solid lines. **D)** Scatter plot showing a significant positive correlation between increase in PCC-DLPFC decoupling during NF-guided meditation (y-axis; session 2 gPPI betas) and decrease in emotional distress from baseline to follow-up (x-axis; DASS-Total difference scores) (FDR-p = 0.018; uncorrected p = 0.003; rho = 0.53; N=32). **E)** Scatter plot showing a non-significant negative correlation between PCC-DLPFC decoupling (y-axis; session 2 gPPI betas) and SMS-Mind slopes from the 1-week meditation practice (x-axis; SMS-Mind slopes) (N = 32). **F)** Scatter plot showing a non-significant positive correlation between PCC-DLPFC decoupling (y-axis; session 2 gPPI betas) and change in BCT accuracy from baseline to follow-up (x-axis; difference in BCT accuracy %) (N = 31). In the scatter plots (D-F), blue dots show experimental group data, red dots show control group data, and grey shaded areas represent 95% confidence intervals around the linear fits (solid black lines). PCC - posterior cingulate cortex; NF - neurofeedback; s1 - session 1; s2 - session 2; EXP - experimental group; CONT - control group; DLPFC - dorsolateral prefrontal cortex; gPPI - generalised psychophysiological; med. - meditation; rest. - restful thinking; FC - functional connectivity; DASS - Depression, Anxiety & Stress Scale; SMS - State Mindfulness Scale; A - anterior; BCT - Breath Counting Task; wk - week; P - posterior; L - left; R - right; I - inferior; S - Superior; * FDR-significant p<0.05; ** FWE- adjusted p<0.05

#### Functional coupling during NF-guided meditation

Next, we examined the effect of NF-guided meditation on the voxel-wise functional coupling of the NF target region (PCC). According to the neurocognitive brain network model of focused attention meditation *(6)*, enhanced breath focus and awareness during meditation are supported by decreased activity in the DMN alongside relative increases in SN and CEN activity. To test this, we assessed group differences in context-dependent functional decoupling (i.e., during meditation vs. rest) in each NF session between the PCC (a DMN hub) and all voxels within the salience network (SN) and CEN, using voxel-wise generalised psychophysiological interactions (gPPI) analysis *(39)*, while controlling for physiological and head motion artefacts, age, sex and sleepiness.

In the first NF session, there were no significant differences in functional decoupling between groups. In the second NF session, we found a significant gPPI cluster (26 voxels) of medium effect size (Cohen’s *d* = 0.59) in the right dorsolateral prefrontal cortex (DLPFC), a core CEN node (FWE-adjusted one-tailed p=0.032; **Figure 4A**; *Table S2)*. In other words, relative to rest, the decoupling between the PCC and right DLPFC was significantly stronger during veridical NF-guided meditation compared to sham NF-guided meditation in the final training session (day 3 in **Figure 2A**).

#### Change in functional coupling from baseline to transfer meditation

The change in functional coupling (Pearson’s correlation functional connectivity (FC)) between the significant gPPI cluster and PCC – from pre-NF baseline meditation to post-NF transfer meditation in the final session with the NF effects – did not significantly differ between groups (p > 0.05; **Figure 4C**).

### Relationship between changes in NF brain target and behavioural outcomes

We sought to determine whether the significant decoupling observed between the PCC and right DLPFC during the final NF session was associated with any of the significant behavioural outcomes. Specifically, we calculated the correlation between the extent of decoupling (gPPI betas) that significantly differed between groups in the final NF session and: (i) changes in mindful awareness during the weeklong meditation practice (SMS-Mind slopes), (ii) changes in emotional distress (DASS-21 difference scores) over the weeklong practice, and (iii) changes in breath counting skill (BCT accuracy differences) from baseline to the end of the weeklong practice. Age and sex effects were partialled out.

We found that the decrease in emotional distress from the week before baseline to the week of post-NF real-world meditation practice was significantly correlated with increase in PCC-DLPFC decoupling during NF-guided meditation (rho(28)=0.53, FDR-adjusted p=0.018; **Figure 4D**). Post-hoc analysis revealed that this association was significant only in the veridical-NF group (rho(12)=0.71, p=0.004), but not in the sham-NF group (rho(12)=0.06, p=0.83).

The PCC-DLPFC decoupling was not significantly correlated with the other behavioural outcomes, i.e., change in mindful awareness (rho(28)=-0.14; p=0.22; **Figure 4E**) or change in breath counting accuracy (rho(27)=0.23; p=0.46; **Figure 4F**). A 5% FDR correction was used to control for multiple comparisons across all PCC-based region-level analyses *(Table S2)*.

## DISCUSSION

In this study, we sought to optimise early meditation practice for well-being by implementing a proof-of-concept 7 Tesla fMRI NF-guided meditation training paradigm. This training involved novices learning to meditate from precise and personalised feedback on PCC activity — a key brain region in the DMN implicated in mental and self-referential processing *(28)*. The paradigm assisted novice meditators in learning focused attention meditation by training them to deactivate their PCC during meditation relative to rest and to understand how PCC activity relates to meditative states.

Our findings revealed that the experimental group, following two consecutive days of contingent NF-guided meditation training, experienced a significantly greater increase in mindful awareness of mental activity during a week of real-world meditation, compared to the control group that received sham NF-guided meditation training. Additionally, the experimental group experienced a significantly greater reduction in emotional distress over this period of post-NF real-world practice. While both groups showed comparable PCC deactivation during NF-guided meditation vs. rest on each training day, the experimental group exhibited significantly greater decoupling between the deactivated PCC and the dorsolateral prefrontal cortex (DLPFC), a region crucial for executive functioning and cognitive control *(40)*, on the final training day. Notably, this increase in PCC-DLPFC decoupling during NF-guided meditation was significantly associated with the decrease in emotional distress over the week of post-NF real-world meditation practice in the experimental group, but not in the control group. Sham NF signals provided to the control group were uncorrelated with their actual PCC signals, yet both groups reported similar perceptions of NF authenticity, task performance, and usefulness of NF for learning meditation, indicating effective blinding and comparable motivation, engagement, expectations, and perception of learning across groups.

The function of the DLPFC, in the CEN, involves cognitive control which includes managing attentional resources to facilitate goal-directed behaviour *(40)* and amplifying attention to target stimuli while filtering out distractions *(41)*. Decoupling between CEN regions (e.g., DLPFC) and DMN regions (e.g., PCC) is thought to release attentional resources from sustained mental processing, facilitating enhanced bottom-up perception of present-moment sensory and bodily stimuli *(10*, *42–45)*. According to the neurocognitive network model of focused attention meditation *(6)*, shifting attention away from default mental processing towards target stimuli, such as breathing sensations, is linked to DMN suppression and relative increases in CEN and SN activity. Furthermore, preliminary evidence suggests that attenuating the DMN relative to CEN through fMRI meditation-NF can positively impact patients with schizophrenia *(30)* and affective disorders *(31)*. Consistent with these functions of DMN-CEN interaction, our NF-guided meditation training, which targeted PCC (DMN) deactivation in healthy novices, significantly enhanced PCC (DMN) -DLPFC (CEN) decoupling during focused attention meditation vs. rest on the final training day. Therefore, compared to the sham NF control group, the experimental group potentially learnt to meditate by exercising greater attentional control and disengagement from mental activity relative to their resting baseline *(46)*.

Sustained meditation practice with effective top-down attentional control has been shown to progressively enhance mindful awareness, facilitating a transition to the more effortless, present-centred, and non-reactive attention observed in experienced meditation practitioners *(7*, *46)*. Early indicators of this process were evident in our study. Proportional to real-world meditation practice time, the experimental group, which demonstrated stronger NF-driven PCC-DLPFC decoupling linked to enhanced top-down control, showed a greater increase in mindful awareness of mental activity during meditation, compared to the control group. Thus, continued self-guided meditation practice, augmented by the insights and attentional enhancements gained from high-precision NF training, may expedite the development of mindful awareness in practice compared to meditating without the benefits of NF-guided training for the same duration *(47)*.

Compared to the control group, the experimental group that meditated with greater mindful awareness of mental activity during the week of real-world meditation practice also experienced greater reduction in negative emotional states during this period, with these reductions correlating with the increased NF PCC-DLPFC decoupling. While prior evidence has shown that a week of meditation practice involving five-minute sessions can mitigate emotional distress in healthy novices *(12*, *48)*, the current study suggests that increased mindful awareness during such five-minute sessions, facilitated by enhanced attentional control acquired from prior high-precision NF training, can further enhance week-long meditation practice by optimising its impact on emotional well-being. Therefore, meditating with greater mindful awareness of self-referential mental processing, can likely facilitate better management of negative emotional states due to quicker recognition of negative thought patterns — a key prognostic factor across stress-related disorders like depression *(49)* — followed by more effective disengagement from these thoughts.

Contrary to our hypothesis, the experimental group showed a significant decline in breath-counting accuracy from baseline to one week post-NF, compared to the control group. We also found the decrease in breath-counting accuracy to be significantly correlated with increase in mindful awareness of mental activity during the week-long practice. Recent evidence demonstrated that the BCT reflects sustained attention more than mindful attention in beginners *(50)*. Therefore, BCT may not reliably capture improvements in focused attention meditation associated with NF training. As such, mindful awareness of and disengagement from mental processes (which includes counting), may interfere with accurately monitoring breath counts during meditation, though further validation of this notion is needed.

Although the experimental group showed significantly stronger PCC-DLPFC decoupling during NF-guided meditation on the final training day, the change in PCC-DLPFC decoupling from pre-NF baseline to post-NF transfer meditation on that day did not differ significantly between groups, due to multiple possible factors. Firstly, an hour-long MRI scanning session with fMRI-NF training can induce substantial mental fatigue *(51)*, potentially hindering transfer meditation performance, particularly for novices who require substantial attentional effort to meditate. Secondly, the effects of fMRI-NF often emerge gradually rather than immediately post-NF *(19)*, which is also observed in our study where significant outcomes emerged over the course of a week. Postponing the transfer task for a follow-up session may enhance sensitivity to neuronal transfer effects.

Previous fMRI meditation-NF studies with small samples of clinical and healthy novices have reported preliminary evidence of increase in state mindfulness *(31*, *34)*, decrease in auditory hallucinations *(30)*, and improvement in attention *(32)* post-NF. However, most of these studies lacked control groups, adjustments for multiple comparisons, real-time physiological denoising, or follow-up assessments, and used continuously updated visual NF (i.e., feedback updated every TR).

In the current paradigm, we included a yoked-sham control group that followed the same strategies and training as the experimental group, except for the feedback contingency, which differed. Both groups deactivated the NF brain target (PCC) similarly and neither group was informed of the existence of a control group. Therefore, control over non-specific factors like motivation, expectancy, reward, task strategy, and engagement was likely well-managed, enabling effective isolation of the effects of learning to meditate through NF target regulation *(52*, *53)*. As a proxy for online physiological correction during NF, we used a control brain region spanning areas typically influenced by physiological responses *(54)*, potentially enabling participants to engage more with mindful attention rather than breath control while learning meditation through NF modulation *(55)*.

Alongside a one-week behavioural follow-up, we incorporated repeated ecological sampling of real-world meditation, which likely increased sensitivity to the evolving benefits of high-precision NF-guided training in natural settings, thereby enhancing the paradigm’s translational value *(20)*. Participants in our study received intermittent feedback after short meditation intervals to address the drawbacks of continuous NF paradigms, which include increases in neurocognitive load during task performance due to simultaneously monitoring the NF signal, managing hemodynamic delays, and processing constantly changing visual inputs *(53*, *56)*. Intermittent NF may have supported novices by minimising distractions associated with continuous NF during meditation, though the impact of intermittent vs. continuous NF on meditation needs further investigation.

In our study, the intermittent feedback, though averaged from meditation blocks, accounted for the voxel-wise, time-varying dynamics of PCC deactivation within each block. The higher spatiotemporal resolution and signal-to-noise ratio of 7 Tesla fMRI *(26*, *27)*, compared to commonly used 3 Tesla fMRI, likely enhanced the reliability of these neuroanatomically focal signal dynamics and in turn the feedback. The specificity of 7 Tesla fMRI also extended to behavioural effects, as the NF training, which targeted the PCC linked to mental processing, specifically enhanced mindful awareness of mental processes but not bodily sensations during subsequent real-world meditation. Finally, we have ensured methodological transparency by adhering to the CRED-nf checklist *(57)*.

The study has limitations that should be noted. The sample size was modest, necessitating cautious interpretation of the findings. The study employed single-blinding which may have introduced experimenter biases. Although participant self-reports indicated successful participant blinding, future research should aim for double-blinded RCTs to robustly validate the efficacy of high-precision NF-guided meditation training. Longer-term follow-ups and more intensive sampling can be valuable, particularly to track enduring linear and non-linear changes in meditation quality, dispositional mindfulness and well-being. Although both groups received identical instructions, potential stagnation or worsening due to sham NF training could have influenced the observed group differences. Future studies could include a no-NF control group in addition to the sham-NF control group to explore whether the observed improvements from veridical NF training extend beyond self-guided, teacher-guided or app-based meditation practice.

Meditation can offer a structured framework with specified instructions for trainees to actively refine NF-acquired skills after the high-precision NF training in real-world settings detached from specialised equipment *(20)*. Our paradigm demonstrates promising evidence that meditation training guided by high-precision NF, compared to meditation training with sham NF, can increase control over disengagement from mental processes, leading to enhanced mindful awareness of mental processes during subsequent real-world meditation practice and greater emotional well-being benefits from this practice. Our findings also underscore the importance of ecological sampling in NF paradigms to advance their translational therapeutic value.

## MATERIALS AND METHODS

### Experimental design

In this single-blind, controlled, longitudinal study, 40 eligible, healthy, adult novice meditators were assigned to either an experimental group (N=20), which meditated with real NF from their own brain activity, or a control group (N=20), which meditated with sham NF (**Table 1**; *Figure S1)*. The participants were blinded to the existence of a control group. The study was approved by the University of Melbourne human research ethics committee (Ethics ID: 24345), and study protocol components are reported in the CRED-nf checklist *(Table S1)*. Novice meditators were defined as having limited or no regular practice, less than 500 hours of lifetime meditation, fewer than 2 retreats, and no retreat experience in the past 2 years. Individuals with a history of neuropsychiatric/neurological conditions, psychoactive/psychedelic substance use, or alcohol-use disorder were excluded. Details are provided in *Supplementary Section 1*.

### Behavioural assessments and self-guided meditation

During the week before baseline, participants completed two five-minute audio-guided focused attention meditation sessions using the mEMA app for familiarisation. At baseline, dispositional mindfulness, mind-wandering, anxiety, and past-month sleep quality were measured (**Table 1**; *Supplementary Section 1)*. Participants completed six 5-minute self-guided focused attention meditation sessions: one at baseline, four at home post-NF training via the mEMA app, and one at the 1-week follow-up. SMS (Mind and Body subscales) *(35)* and SSS *(37)* were measured before and after each session. Pre-meditation SMS assessed mindful awareness during 5 minutes before meditation, while post-meditation SMS assessed it during the 5-minute meditation. The DASS-21 *(36)* measured negative emotional states (related to depression, stress, anxiety) at baseline and 1-week follow-up, reflecting emotional distress during the week prior to baseline and the post-NF week of self-guided meditation. The BCT *(38)*, an objective mindfulness proxy, was assessed at baseline and follow-up, measuring task (% of correct breath count cycles) and probe (% of correct breath counts reported when probed) accuracy.

### MRI data acquisition

MRI data was acquired on a 7 Tesla MRI scanner (Siemens Magnetom 7T plus) using an 8/32 PTX/RX channel head coil. High-resolution whole-brain T1-weighted (T1w) anatomical images (3D-MP2RAGE; 0.75mm isometric voxel size; TE/TR=2ms/5000ms) and functional images (1.6mm isometric voxel size; TE/TR=22ms/800ms; multiband acceleration=6) were acquired using multiband gradient-echo echo-planar imaging (EPI) sequence *(Supplementary Section 1)*. Concurrent physiological measurements were acquired using MRI-compatible respiration belt and a pulse oximetry sensor, with data from the latter excluded due to faulty recordings.

The fMRI NF-guided meditation training was conducted over two consecutive days for each participant. Each fMRI session (**Figure 2B**) comprised a baseline meditation task (2.5 minutes) without NF, three NF-guided meditation runs (10 minutes each), and a transfer meditation task (2.5 minutes) without NF. Each NF run had a blocked design: three blocks of rest (51s each) and six blocks of focused attention meditation (26s each). NF scores were visually displayed *(Figure S2)* after each meditation block, with three pairs of meditation and NF blocks between each pair of rest blocks. After each fMRI run, SSS sleepiness ratings were acquired. After each fMRI session, participants rated the utility of NF, their meditation performance, and correspondence between NF scores and their meditative attention using 5-point Likert scales, and summarised their in-scanner meditation and rest strategies.

### Real-time fMRI

Real-time fMRI preprocessing and analysis were conducted using Turbo-BrainVoyager (v4.2) and MATLAB, with visual cues and feedback managed by Psychtoolbox (v3.1). Preprocessing steps included coregistration to anatomical and MNI space, motion correction, spatial smoothing, linear detrending, and physiological control through nuisance regression using BOLD PSC from a confound ROI *(Figure S4A)* linked to physiological artefacts *(Supplementary Section 1)*.

The target ROI for NF used in this study was the bilateral ventral PCC (**Figure 3A**), defined using the Schaefer brain atlas *(58)* in standard MNI space. Real-time voxel-wise PSC of the target PCC ROI was estimated through incremental GLM in Turbo-BrainVoyager. The mean PSC from each timepoint was estimated from the 33% of PCC voxels most responsive to meditation vs. rest, enabling dynamic PCC personalisation. Real-time physiological control was implemented in MATLAB using cumulative GLM, with PCC PSC as the response variable and confound ROI PSC as the predictor. The residualised PCC PSC was averaged within each meditation condition (26 seconds) to estimate the NF score for feedback displayed after the meditation condition *(Supplementary Section 1)*. The scores were visualised on a thermometer bar with 20 levels, where higher levels indicated more negative PCC PSC (i.e., deactivation relative to rest) linked to greater meditative focus.

### Statistical Analyses

#### Offline MRI preprocessing and analysis

All MRI data was preprocessed using fMRIPrep (v23.2.1) *(Supplementary Section 1)*. FMRI preprocessing steps included spatial inhomogeneity distortion correction, non-linear coregistration to anatomical and MNI space, head motion correction, and spatial smoothing with a Gaussian kernel of 2mm full-width half-maximum (FWHM). Nuisance regressors for BOLD fMRI denoising included 24 head motion parameters, top five aCompCor parameters for physiological noise, cosine regressors for high-pass filtering, nine RETROICOR respiration correction regressors, and regressors for non-steady state magnetization effects from initial fMRI volumes. After fMRIprep quality checks, fMRI data from six participants were discarded due to reduced BOLD signal quality, leaving data from 34 participants (17 experimental, 17 control) for MRI-related analysis.

Subject-level GLMs were used to model voxel-wise BOLD responses to meditation, rest, cue and feedback conditions during each NF run, while controlling for variance associated with the nuisance regressors. Second-level GLM was used to quantify the mean PCC BOLD responses across NF runs for the meditation vs. rest contrast, while controlling for mean framewise displacement (mFD). Group-level differences in mean PCC activation during meditation vs. rest, with age, sex and average SSS ratings as covariates were assessed using a third-level GLM for each fMRI session. Similar procedure was followed for the gPPI analysis examining group differences in functional decoupling between PCC and all voxels in the DMN and CEN, in line with the neurocognitive network model *(6)*. The first-level model included NF task predictors, PPI predictors (PCC BOLD time course × condition), and PCC BOLD time course, with nuisance regressors. Group differences per NF session were identified using non-parametric permutation testing (10,000 permutations, uncorrected p<0.005 cluster-forming, FWE-corrected p<0.05 across clusters). Functional decoupling between PCC and significant NF gPPI cluster/(s) during baseline and transfer tasks was assessed using Pearson’s correlation functional connectivity (FC), and group differences in FC change scores (transfer minus baseline) were analysed with GLM (ANCOVA), controlling for mean SSS, mFD, age, and sex.

### Behavioural data analysis

Behavioural data from two participants was excluded due to doubtful compliance with at-home app usage, resulting in N=38 participants (19 per group) for behavioural analyses. A GLM (ANCOVA) was used to detect group differences in the total DASS-21 difference scores (follow-up minus baseline), with age and gender as covariates.

To isolate the change in mindful awareness during real-world meditation practice, pre-meditation SMS scores, mean SSS ratings, age and sex were regressed out from the post-meditation SMS scores of each 5-minute meditation session. Repeated measures ANCOVAs were performed on the resulting residuals from each participant across the six meditation sessions (timepoints), with time since baseline as the predictor variable. Each ANCOVA produced one regression slope per participant for each SMS subscale, which was subsequently entered into one-way ANOVA to examine group differences per subscale.

Brain-behaviour association was examined through Pearson’s partial correlation between each significant behavioural outcome (here, DASS-21, SMS slopes, and BCT accuracy) and significant NF target changes (here, PCC-seeded gPPI cluster), controlling for age and sex. Group differences in post-NF Likert-scale self-ratings were analysed using Wilcoxon rank sum tests, while correlations between online and offline PCC signals, as well as actual and sham PCC signals from the control group, were assessed using Pearson’s correlation. Where applicable, t-statistics were transformed to Cohen’s *d* effect sizes. Multiple comparisons were controlled using FDR with a significance threshold of p<0.05. Detailed description of methods is provided in *Supplementary Section 1*.

## Acknowledgements

The authors thank the National Imaging Facility at the Melbourne Brain Centre Imaging Unit for access to the 7 Tesla MRI scanner, and technical support from Rebecca Glarin, Braden Thai, and Tudor Sava; Siemens for MRI pulse sequences; University of Melbourne IT support (Kire Kalajdziovski) and the Turbo BrainVoyager team (Dr. Michael Lührs & team) for assistance in establishing the neurofeedback facility; and the University of Melbourne’s Research Computing Services for the Spartan high-performance computing platform. Special thanks to all the volunteer participants for their time and effort.

## Data and Code Availability

Upon request and necessary ethics approvals from the requester, data will be deidentified and shared. Code used for real-time fMRI-NF, and offline data analysis can be accessed via the project’s Github repository (https://github.com/saampras/Melbourne-7T-fMRI-Neurofeedback-guided-meditation/tree/main).

