## Supplementary Material for "High-precision neurofeedback-guided meditation training optimises real-world self-guided meditation practice for well-being"

This PDF file includes

### Supplementary Section 1: Methods

#### Sample

51 healthy, adult volunteers were found eligible and willing to participate, of which 11 did not complete the study requirements for the following reasons: withdrawal after the baseline session (N=2), withdrawal during MRI scanning due to discomfort (N=7), and cancellations caused by MRI scanner malfunctioning (N=2). The final sample consisted of 40 participants (**Figure S1** for CONSORT schematic).

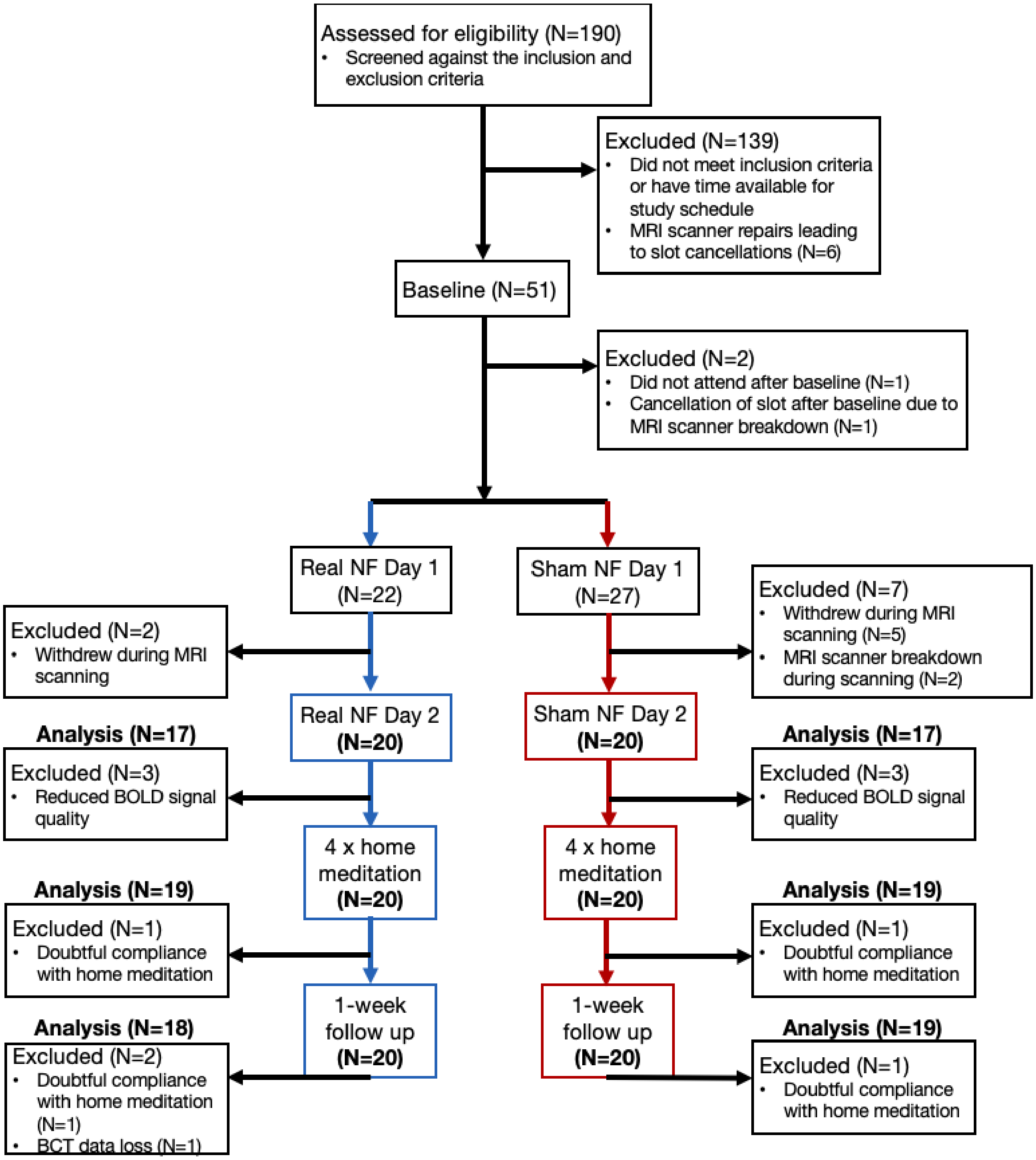

**Figure S1:** CONSORT schematic illustrating participant recruitment, dropout, completion, and data inclusion for analysis.

Volunteers were recruited via email advertisements from the local community and provided written informed consent to participate. The 40 participants were assigned to either an experimental group (N=20), which meditated with real NF from their own brain activity, or a control group (N=20), which meditated with sham NF derived from the brain activity of an experimental participant with similar meditation experience (i.e., yoked-sham NF). The participants were not informed about the existence of a control group in the study, and were consequently blinded to their group assignment. Every participant received expert guidance and instruction on meditation technique, and a 3D printed model of their high-resolution anatomical brain image as reward for volunteering.

The inclusion criteria were: age between 19 and 50 years; self-reported interest in learning meditation; fluency in English; beginner at meditation - broadly defined as having irregular, sparse or no meditation practice, no meditation retreat experience over the past 2 years, cumulative lifetime meditation experience under 500 hours and lifetime participation in less than 2 meditation retreats; and completed the two prescribed audio-guided meditation sessions prior to baseline assessments. The exclusion criteria were: self-reported lifetime clinical diagnoses of any neuropsychiatric (e.g., psychosis, addictions, depression, anxiety) or neurological (e.g., traumatic brain injury, epilepsy) disorders; lifetime consumption of any psychoactive medication (e.g., antidepressants, benzodiazepines, anti-psychotics) with or without prescription; regular or recent consumption of psychoactive (e.g., cannabis) or psychedelic (e.g., Psilocybin) drugs; alcohol-use disorder with a score of $\geq$ 4 for males and $\geq$ 3 for females in the Alcohol Use Disorders Identification Test (AUDIT-C) (*59*); or contraindications to MRI scanning. The CRED-nf checklist indicating the study protocol components is presented in **Table S1**.

**Table S1**: Consensus on the Reporting and Experimental Design of clinical and cognitive-behavioural Neurofeedback studies (CRED-nf) best practices checklist (*57*)

| Domain | Item no. | Checklist item | Reported on page no. in main manuscript |
|---|---|---|---|
| **Pre-experiment** | | | |
| | 1a | Pre-register experimental protocol and planned analyses | N/A - proof-of-concept study |
| | 1b | Justify sample size | N/A - proof-of-concept study |
| **Control groups** | | | |
| | 2a | Employ control group(s) or control condition(s) | 4,5,6,7 |
| | 2b | When leveraging experimental designs where a double-blind is possible, use a double-blind | N/A - proof-of-concept study was single-blind |
| | 2c | Blind those who rate the outcomes, and when possible, the statisticians involved | N/A |

| | | | |
|---|---|---|---|
| | 2d | Examine to what extent participants and experimenters remain blinded | 8,22 |
| | 2e | In clinical efficacy studies, employ a standard-of-care intervention group as a benchmark for improvement | N/A - proof-of-concept study in healthy adults |
| **Control measures** | | | |
| | 3a | Collect data on psychosocial factors | 4,6,7,8 |
| | 3b | Report whether participants were provided with a strategy | 6,8,14 |
| | 3c | Report the strategies participants used | 8,20 |
| | 3d | Report methods used for online-data processing and artefact correction | 20,21 |
| | 3e | Report condition and group effects for artefacts | 8 |
| **Feedback specifications** | | | |
| | 4a | Report how the online-feature extraction was defined | 20 |
| | 4b | Report and justify the reinforcement schedule | 7,17,20 |
| | 4c | Report the feedback modality and content | 6,7,20 |
| | 4d | Collect and report all brain activity variable(s) and/or contrasts used for feedback, as displayed to experimental participants | 11,12,13,20,21 |
| | 4e | Report the hardware and software used | 19,20 |
| **Outcome measures** | | | |
| Brain | 5a | Report neurofeedback regulation success based on the feedback signal | 11,13 |
| | 5b | Plot within-session and between-session regulation blocks of feedback variable(s), as well as pre-to-post resting baselines or contrasts | 13 |
| | 5c | Statistically compare the experimental condition/group to the control condition(s)/group(s) (not only each group to baseline measures) | 11,12,13,21 |
| Behaviour | 6a | Include measures of clinical or behavioural significance, defined a priori, and describe whether they were reached | 8,9,10,19,21,22 |

| | | | |
|---|---|---|---|
| | 6b | Run correlational analyses between regulation success and behavioural outcomes | 12,13,22 |
| **Data storage** | | | |
| | 7a | Upload all materials, analysis scripts, code, and raw data used for analyses, as well as final values, to an open access data repository, when feasible | Data will be shared upon request and satisfaction of required ethics approvals. Data will also be shared with global consortia such as ENIGMA-Meditation and ENIGMA-Neurofeedback. |

Darker shaded boxes represent *Essential* checklist items; lightly shaded boxes represent *Encouraged* checklist items.

### Baseline assessments

At baseline, dispositional mindfulness, dispositional mind-wandering, dispositional anxiety, and sleep quality in the past month were measured using Five Facet Mindfulness Questionnaire (FFMQ) (*60*), Mind-Wandering Questionnaire (MWQ) (*61*), State and Trait Anxiety Inventory, Trait module (STAI-T) (*62*), and Pittsburgh Sleep Quality Index (PSQI) (*63*) respectively. These baseline assessments facilitated the detection of any significant pre-existing differences between groups before the study commenced.

#### Five Facet Mindfulness Questionnaire (FFMQ)

FFMQ (*60*) was used to measure dispositional mindfulness, i.e., general tendency to be mindful in daily life. FFMQ shows high internal reliability (Cronbach's $\alpha$ > 0.85) (*64*), and comprises 39 self-report questions covering 5 mindfulness facets, i.e., observing, describing, acting with awareness, non-reactivity to inner experiences, and non-judging of inner experiences. Questions are scored from 1 to 5, with higher scores indicating greater tendency towards mindfulness in daily life.

#### Mind-Wandering Questionnaire (MWQ)

MWQ (*61*) was used to assess dispositional mind-wandering. It consists of six statements, each self-rated on a 6-point Likert scale. MWQ demonstrates high internal reliability (Cronbach's $\alpha$ = 0.85) (*61*), and evaluates the extent of task-unrelated thoughts in daily activities, capturing both deliberate and spontaneous mind-wandering. Lower MWQ scores signify lower levels of dispositional mind-wandering, which are typically also related to higher mindful awareness in daily life (*61*).

##### State and Trait Anxiety Inventory, Trait module (STAI-T)

STAI-T (*62*) was administered to measure dispositional anxiety. STAI-T comprises 20 self-report questions with 4-point Likert scale responses. Lower scores indicate lower levels of dispositional anxiety. STAI-T shows high internal reliability (Cronbach's $\alpha$ > 0.85) and has been validated to assess anxiety in research and clinical settings (*65*). Greater tendencies towards mindful qualities have been associated with lowered levels of dispositional anxiety (*66*).

##### Pittsburgh Sleep Quality Index (PSQI)

PSQI (*63*) was used to evaluate sleep quality of participants one month prior to study commencement. The PSQI is a validated and widely-used clinical instrument with Cronbach's $\alpha$ > 0.70 (*67*). It comprises 19 individual items spanning seven components: subjective sleep quality, sleep latency, sleep duration, habitual sleep efficiency, sleep disturbances, use of sleeping medication, and daytime dysfunction. Lower total PSQI scores signify higher overall sleep quality over the past month. PSQI enabled identifying between-group differences in baseline sleep quality, which can potentially influence subsequent meditation and mindfulness states (*68*).

#### Outcome assessments

##### Depression, Anxiety and Stress scale (DASS-21)

Meditation and mindfulness training can alleviate distress in healthy individuals (*69*). DASS-21 (*36*) was used to quantify emotional distress over a 1-week time-interval. DASS-21 is a validated self-report questionnaire with high internal reliability (Cronbach's $\alpha$ > 0.8) in non-clinical samples (*70*). It comprises 21 questions with 4-point Likert rating responses, providing a quantitative measure of distress along the 3 axes of depression, anxiety and stress over the past week. The total DASS-21 score (Cronbach's $\alpha$ = 0.93) represents overall emotional distress, serving as a composite measure of negative emotional states related to depression, stress, and anxiety. Lower scores signify lesser distress.

##### State Mindfulness Scale (SMS) & Stanford Sleepiness Scale (SSS)

The SMS (*35*) is a self-report tool that measures perceived attention and mindful awareness of present-moment experiences over a specified period and context. When the context involves meditation, SMS responses capture mindful awareness during meditation.

SMS comprises 21 statements with 5-point ratings divided into two subscales based on the object of mindful attention and awareness, i.e, SMS Body (Cronbach's $\alpha$ > 0.80) for bodily sensations (6 statements), and SMS Mind (Cronbach's $\alpha$ > 0.90) for mental events (e.g., emotions, thought patterns; 15 statements). Higher scores indicate greater mindful awareness. SMS has consistently demonstrated excellent internal reliability (Cronbach's $\alpha$ range = 0.80 to 0.95)(*71*). Crucially, SMS enables capturing dynamic changes in mindful awareness associated with meditation training, practice and expertise(*71*).

Alongside each SMS measurement, momentary levels of subjective sleepiness were recorded using the SSS (*37*). The SSS, a widely used sleepiness measurement tool, consists of a single 8-point Likert scale where lower ratings indicate greater alertness. SSS ratings were used to control for the impact of drowsiness on meditation states and awareness.

##### Breath Counting Task (BCT)

The Breath Counting Task (BCT) (*38*), an objective proxy of mindfulness skill, was used to evaluate attention to breathing cycles and their counts. In this task, participants count their breaths from 1 to 9 cyclically, registering each count with a button press. A left arrow button was pressed for the first eight counts, while a right arrow button was pressed for the ninth count of each cycle. At five pseudo-random instances during the task, participants were probed to verbally report their current breath count.

Task accuracy is assessed by the percentage of correct count cycles, and probe accuracy is determined by the percentage of correctly reported counts when probed. The physiological accuracy of the breath counts was evaluated by correlating the count rates with the breathing rates recorded using a respiration belt (Vernier Science Education, Oregon, USA) worn during the task.

#### Study Procedure

The complete study design is illustrated in **Figure 1A** (main text). The study included several stages: baseline assessments and a self-guided meditation session, two consecutive days of fMRI NF-guided meditation training, four self-guided meditation sessions at home during the following week, and follow-up assessments with a self-guided meditation session after one week.

#### Pre-baseline

During the week before baseline measurements, participants completed two five-minute audio-guided sessions of focused attention meditation provided by the Epworth Clinic (https://www.epworth.org.au/our-services/mental-health/resources). These sessions familiarised participants with the meditation technique. Participants used the mobile Ecological Momentary Assessment (mEMA) app (https://ilumivu.com/solutions/mobile-health/) on their phones, completing one session per day at home. The mEMA approach captures real-world experiences closer to when and where they occur, reducing response biases and memory distortions.

#### Baseline with self-guided meditation (Day 1)

On day 1, participants completed various assessments: FFMQ for dispositional mindfulness, MWQ for dispositional mind-wandering, STAI-T for dispositional anxiety, PSQI for 1-month sleep quality, BCT for breath counting skill, and DASS-21 for 1-week emotional distress. They then engaged in a 5-minute session of self-guided, silent, eyes-closed meditation with focused attention on their breath, following instructions adapted from literature (*72*)**.**

The instructions were as follows:
*“Sit comfortably and upright. Close your eyes and focus on the actual sensations of breath entering and leaving the body. There is no need to think about the breath, no need to change or control it. Just experience the sensations of it as you naturally breathe in and out. Whenever you notice that your awareness is no longer on the breath and you have been distracted by thoughts, emotions or other sensations, gently bring your awareness back to the sensations of breathing. You will hear a bell ring to mark the start and end of your session.”*

Before and after the meditation, participants completed the SMS and SSS. The pre-meditation SMS measured mindful awareness during the 5 minutes before the session, while the post-meditation SMS measured mindful awareness during the meditation period.

#### fMRI neurofeedback (NF) - guided meditation training (Day 2 & Day 3)

One day after the baseline, participants began fMRI NF-guided meditation training, which included two sessions over consecutive days (one session per day). NF-guidance for meditation was based on BOLD activity in the posterior cingulate cortex (PCC).

#### Self-guided meditation at home (during week after NF)

During the week after NF-guided meditation training, participants used the mEMA app at home to complete four 5-minute sessions of self-guided, silent, and eyes-closed focused attention meditation, while applying insights from their NF training.

The instructions were as follows:

"

1. *Please find a comfortable place and posture.*
2. *You will hear a bell ring in the audio to mark the start and end of your session.*
3. *You may use earphones/headphones for this session. If using, please connect it to your phone before proceeding.*
4. *To meditate, you will follow the same basic technique you practised during the fMRI neurofeedback training.*
5. *Sit comfortably and upright.*
6. *Focus on the sensations you feel in your abdomen (belly and chest area) as you breathe normally.*
7. *Try not to take deeper or shallower breaths than usual.*
8. *Whenever you notice your mind being distracted by thoughts, emotions or other sensations, gently acknowledge and bring your focus back to the abdomen.*
9. *You can keep your eyes closed during the session.*
10. *Start meditating when you hear the bell in the audio. You can stop meditating when you hear the next bell after 5 minutes.*

"

Although participants were advised to focus on breathing sensations at the abdomen (similar to the NF sessions), they were also free to instead direct their attention to other areas of the body associated with breathing sensations if those felt more convenient. Before and after each session, participants completed the SMS and SSS questionnaires to measure mindful awareness before and during meditation. They were required to complete four sessions, with no more than one per day, and could schedule them flexibly. Text reminders were sent to promote adherence.

#### Follow up with self-guided meditation (1 week after NF)

Approximately one week after the NF training, participants visited the laboratory for the final 5-minute session of self-guided, silent, and eyes-closed meditation (following same instructions as above), with SMS and SSS administered before and after. They were also administered the DASS-21, to examine emotional distress over the past week, and the BCT, to evaluate breath counting skill.

Some secondary measures were also acquired at different time points, including pre- and post-NF SMS, go/no-go task and working memory task, which will be reported separately.

#### MRI data acquisition

MRI data was acquired on a 7 Tesla MRI scanner (Siemens Magnetom 7T plus) at the Melbourne Brain Centre Imaging Unit (MBCIU) using an 8/32 PTX/RX channel head coil. High-resolution T1-weighted (T1w) anatomical images (3D-MP2RAGE; 0.75mm×0.75mm×0.75mm; TE/TR=2ms/5000ms) were denoised and corrected for Radio Frequency (RF) inhomogeneity. Whole-brain functional images (1.6mm×1.6mm×1.6mm; TE/TR=22ms/800ms; multiband acceleration=6; field-of-view=208mm; matrix size=130×130; 84 slices; slice thickness=1.6mm; flip angle=$45^0$; P-A phase encoded) were acquired every 800 ms using a multiband gradient-echo echo-planar imaging (EPI) sequence. Concurrent physiological measurements were acquired using Siemens MRI compatible abdomen respiration belt and pulse oximetry finger sensor. The pulse oximetry data was however excluded due to faulty recordings. An MRI compatible two-button response box was used for participants to provide behavioural ratings inside the scanner.

#### Real-time fMRI

##### Acquisition

The Turbo-BrainVoyager software (version 4.2; Brain Innovation, Maastricht, the Netherlands), installed on a dedicated computer, was used for real-time fMRI processing. MATLAB Psychtoolbox (version 3.1), installed on a separate computer, managed the timed visual display of cues, instructions, and feedback presentation during MRI scanning. The Siemens MRI console computer received the reconstructed MRI DICOMs. All three computers - the one running Turbo-BrainVoyager, the one using Psychtoolbox, and the MRI console computer - were connected to a common network via Transmission Control Protocol (TCP). Reconstructed MRI DICOM images were exported from the MRI console computer via the Siemens IDEA command tool to Turbo-BrainVoyager in real-time. NF scores were calculated and visualised in the computer running MATLAB and Psychtoolbox, using outputs received in real-time from Turbo-BrainVoyager.

#### Preprocessing

Following anatomical scan and prior to fMRI scanning, skull stripping and brain image extraction were performed on the anatomical image in Turbo-BrainVoyager. Real-time fMRI preprocessing in Turbo-BrainVoyager included: linear coregistration of fMRI reference volume (run's first volume) with the anatomical and subsequently the Montreal Neurological Institute (MNI) standard stereotactic space, motion correction with 3D trilinear interpolation (realigning each fMRI volume to reference volume), and spatial smoothing (Gaussian kernel of 3.2 mm full width at half maximum). Additionally, real-time voxel-wise linear detrending and 6-parameter head motion regression were performed through incremental general linear modelling (iGLM) to correct for linear and non-linear confounds affecting the BOLD signal. Real-time physiological control was also implemented through cumulative GLM (cGLM) regression with the BOLD signal from a confound region-of-interest (ROI) in MATLAB.

#### Analysis

The target ROI for NF used in this study, i.e., bilateral ventral PCC (**Figure 3A** - main text), was defined using the Schaefer brain atlas (*58*) in standard MNI space. The active task involved focused attention meditation, while the control task involved rest, with NF scores calculated intermittently after each meditation condition.

The voxel-wise percent signal change (PSC) of the PCC was estimated in real-time using incremental general linear modelling (iGLM) in Turbo-BrainVoyager. Residuals from incrementally regressing out nuisance regressors served as the detrended baseline, providing a stable estimate of the PCC BOLD signal during the rest condition. At each timepoint, the mean PSC was estimated from the 33% of PCC voxels most responsive to meditation vs. rest in iGLM, enabling personalised NF and accommodating changes in ROI engagement over time and with learning.

PSC from a proxy confound ROI (shown in **Figure S4A**) (including midline anterior and posterior grey matter, white matter, and ventricular structures), which is generally prone to physiological artifacts (*54*), was also estimated. Real-time physiological control was implemented in MATLAB using a cGLM, with PCC PSC as the response variable and control ROI PSC as the predictor. The residualised PCC was used to estimate the NF score in MATLAB after each meditation condition.

##### Feedback Calculations

For each meditation condition (32 TRs long), the NF score (ns) was calculated from the mean residual PSC values of the PCC (p) across timepoints (TRs) within that condition, excluding the initial 12 TRs (9.6 seconds). Specifically, seven initial TRs were excluded to account for hemodynamic lag, and five initial TRs were excluded to account for transitioning into the meditative condition. The score was then derived from the mean of the remaining 20 TRs, using a standard PSC threshold (thr) of -2%, on an integer scale of 1-20. The neurofeedback score for each meditation condition was estimated in MATLAB as follows:

$$ns = (((\sum_{p=t+e+1}^{t+n} p)/(n-12)) / thr) \times 20 ,$$

Where,

*e* denotes the number of initial timepoints within the meditation condition that were excluded during calculations,

*n* denotes the total number of timepoints within the meditation condition,

*t* denotes the total number of timepoints up to the start of the meditation condition.

Negative scores were adjusted to 1, and positive scores greater than 20 were capped at 20. This ensured an increase in NF scores exclusively reflected negative PSC (i.e., deactivation) in the PCC during meditation compared to resting baseline.

#### fMRI NF-guided meditation training procedure

Participants completed two identical fMRI NF-guided meditation training sessions over consecutive days. They were instructed to avoid caffeine for two hours before each session.

Prior to the first MRI session, participants were introduced to the meditation and control conditions, as well as the visual NF interface, in a mock scanner. They were thoroughly briefed on the NF paradigm and meditation technique to ensure adequate familiarisation.

The debriefing instructions prior to entering the scanner were as follows:

“

1. *Meditation with eyes open lying inside the MRI scanner can be difficult for some people. But training to focus attention on your abdomen sensations despite this difficulty can potentially strengthen your attention and meditation skills.*
2. *The feedback score represents your average level of focus and awareness of breathing sensations. This means that consistently good focused attention and awareness throughout a meditation period is likely associated with higher scores, compared to short instances of good or very good focused attention and awareness.*
3. *Try to use the MRI training to learn and become more aware of these associations from direct experience of meditation.*
4. *Use the feedback display from each meditation period as a guidance to really understand and evaluate what it feels like to be optimally focused and aware, and not optimally focused and aware. You can then try to apply this insight during meditation without neurofeedback.*
5. *During rest periods, do not meditate. You can think about whatever comes to mind. Some examples: planning your day/week; thinking about your memories, family, friends, etc.; thinking about adjectives that describe your personality; etc.*
6. *During meditation periods, focus and observe the sensations and feeling of breathing, for example at the abdomen (part in contact with the belt). Examples of sensations include tightness, warmth, coolness, tingling, touch, etc.*
7. *During meditation, the quality of focus and awareness is higher when distractions are lower. Distractions include thinking, day-dreaming, stories, imagination, narratives, interpretations, etc.*
8. *Please try to keep your head as still as possible during each scanning session. Head movements can negatively impact MRI signal, and sometimes affect your feedback scores.*
9. *Go in with an open mind and be patient.*

*“*

Participants were advised to focus on breathing sensations at the abdomen, as it could have facilitated a clear perception of sensations due to the physical contact of the respiration belt. However, they were also encouraged to instead direct their attention to other areas of the body associated with breathing sensations if those felt more convenient.

During the anatomical scan before fMRI measurements, participants had another practice NF run for familiarisation. The first fMRI run involved a baseline meditation task without NF (self-guided meditation) for 2.5 minutes, with instructions to keep their eyes open and attend to breathing sensations in the abdomen area while breathing normally.

The instructions provided during the fMRI sessions inside the scanner were as follows:
*Baseline meditation task (no NF):*
*"Please keep your eyes open throughout the session. You will NOT receive neural feedback for this session. To MEDITATE: Focus on the sensations and feelings in your stomach area while you breath normally as usual. There is NO need to think about the sensations or change it. Just experience the sensations and feelings as they happen. Whenever you notice that your awareness is no longer on the stomach area, gently bring your awareness and focus back to the sensations of your stomach area. When you see a grey cross on the screen, keep your eyes fixed on the cross and start meditation. Breathe normally as usual."*

*Meditation condition within NF runs*
*"To MEDITATE: Keep your eyes open. FOCUS on the sensations in your stomach area when the cross appears. Breathe normally as usual. You will see feedback on your performance after each meditation period."*

*Rest condition within NF runs*
*"Keep your eyes open. THINK whatever comes to mind freely, when the cross appears. Breathe normally as usual."*

*Transfer meditation task (no NF)*
*"Please keep your eyes open throughout the session. You will NOT receive neural feedback for this session. You will meditate by applying what you have learnt during the neurofeedback training so far. The technique of meditation is the same. When you see a grey cross on the screen, keep your eyes fixed on the cross and start meditation. Breathe normally as usual."*

After the baseline meditation run, participants completed three identical fMRI NF runs (710 TRs each) with a blocked design: three blocks of rest (51s each) and six blocks of focused attention meditation (26s each) (**Figure 1B** - main text). Visual NF was displayed intermittently after each meditation block, showing the average PCC PSC from the most recent block and a history of previous scores (**Figure S2**). In each NF run, there were three pairs of meditation and NF blocks between each pair of rest blocks. Short 26-second meditation blocks were used based on previous evidence that they are suitable for novice meditators in fMRI tasks (*8*).

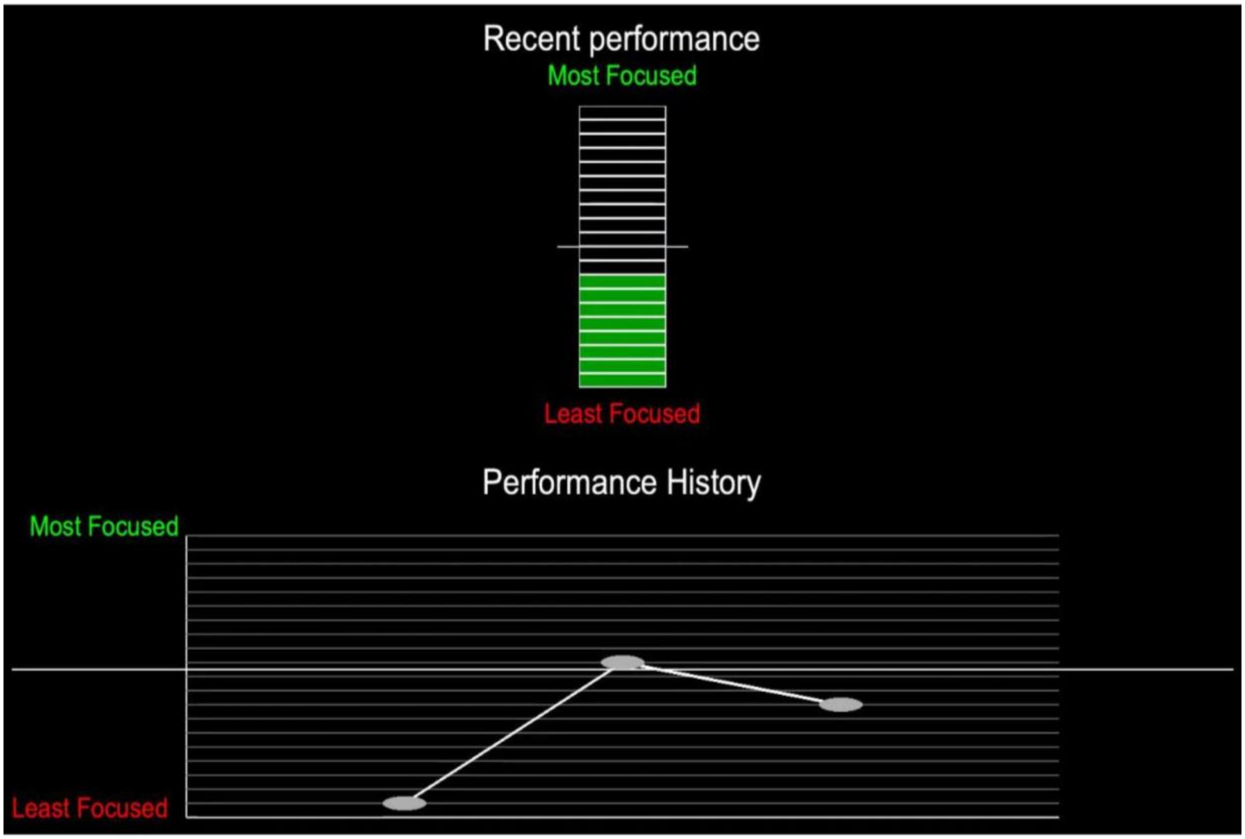

***Figure S2*** *- The visual NF display shown to participants intermittently during the fMRI NF-guided meditation training. Presented for approximately 10 seconds at the end of each meditation block, the display featured a bar under 'recent performance' representing the NF score from the recently concluded meditation block. Participants were instructed to fill up the bar through focused attention meditation. Under 'performance history', a line graph displayed scores from the last two meditation conditions alongside the current one, offering participants context on their performance relative to recent trials.*

The final fMRI run, after the NF runs, involved a transfer meditation task identical to the baseline meditation task, without NF, for 2.5 minutes (188 TRs). After each fMRI run, participants used buttons to rate their current level of sleepiness via the SSS. Participants kept their eyes open throughout the fMRI sessions.

Throughout the NF training, participants were instructed to maintain normal breathing during meditation to minimise confounding effects of breathing variations that can negatively affect meditation fMRI studies (*6*, *8*). Such variations in breathing could otherwise encourage and teach participants to employ breath control instead of the intended meditation technique to increase their NF scores. MRI meditation task instructions were adapted from (*8*). After each fMRI session, participants reflected on the utility of NF training, their meditation performance, and the correspondence between NF scores and their meditative attention using 5-point Likert scales, and provided written summaries of their strategies for the meditation and rest conditions during the NF runs.

#### Offline MRI preprocessing

All MRI data was converted to BIDS format (v1.4.0) using the dcm2niix tool (version 1.0.20230411). Preprocessing was implemented in fMRIPrep 23.2.1.

For the T1w anatomical images, intensity non-uniformity correction and skull stripping were performed using Advanced Normalisation Tools (ANTs) (*73*). FSL FAST was used to segment the brain-extracted T1w images into cerebrospinal fluid (CSF), white matter (WM) and grey matter (GM) (*74*). For each participant, the brain-extracted T1w images from the two MRI sessions were merged into a single unbiased T1w image (equidistant from both the source T1w images) using FreeSurfer's mri_robust_template (*75*). The merged T1w images were spatially normalised to the standard space (MNI152NLin2009cAsym) using ANTs volume-based non-linear registration.

For each fMRI run per subject, the following preprocessing steps were performed in fMRIPrep (*76*). An EPI BOLD reference volume was generated. Spatial distortions in the EPI BOLD images caused by B0 magnetic field inhomogeneities were corrected by estimating fieldmaps using FSL Topup (*77*), utilising opposite phase-encoded (P-A and A-P) spin-echo EPI images acquired during each fMRI session. The estimated fieldmaps were then aligned with the EPI BOLD reference volume using rigid-registration. The distortion-corrected EPI BOLD reference was co-registered to the T1w reference using boundary-based registration with six degrees of freedom in FreeSurfer (*78*). Head-motion parameters relative to the EPI BOLD reference, including transformation matrices and six rotation and translation parameters, were estimated before spatiotemporal filtering using FSL mcflirt (*79*). All resampling was done with a single interpolation step, resulting in preprocessed, spatially normalised fMRI BOLD data. Spatial smoothing was then applied to the preprocessed data using a Gaussian kernel with a 2 mm full-width half-maximum (FWHM) in FSL.

The smoothed preprocessed BOLD data was denoised using nuisance regressors. Specifically, the nuisance regressors included 24 parameters for head motion correction (six head motion parameters, their derivatives and squared derivatives), the top five anatomical component-based noise correction (aCompCor) parameters for physiological noise correction (*80*), cosine regressors for high-pass temporal filtering, nine regressors for respiration correction (including one for respiratory volume change) estimated from respiration belt data using RETROspective Image CORrection (RETROICOR) within the PhysIO toolbox (*81*), and regressors for the initial fMRI volumes with non-steady state magnetization effects.

Following quality control (QC) checks of the fMRIprep preprocessing reports, fMRI data from six participants were discarded due to reduced BOLD signal quality. All subsequent MRI-related analyses were conducted on a sample of 34 participants (N=17 experimental, N=17 control).

#### Offline behavioural data analysis

##### Change in emotional distress

We measured DASS-21 at follow-up to evaluate emotional distress during the 1-week self-guided meditation period after NF training, compared to the week before baseline. Difference scores (follow-up minus baseline) for total DASS-21 were calculated, and an Analysis of Covariance (ANCOVA) model within the General Linear Model (GLM) framework was used to detect significant group differences in these difference scores, with age and gender as covariates. The strong correlations between the DASS-21 subscales at both timepoints (rhos (36)>0.79, p<0.0001) justified the use of total DASS-21 score instead of subscale scores for this analysis. In each group, one participant demonstrated doubtful compliance with the app usage at home, resulting in data exclusion and N=38 participants (N=19 per group) for the behavioural analyses.

##### Change in mindful awareness during real-world meditation

We quantified the change in trajectory (i.e., slope) of mindful awareness during self-guided meditation using SMS scores collected immediately after each of the six five-minute sessions. The first meditation session was conducted at baseline before NF training.

To account for non-specific fluctuations in transient baseline mindful awareness before meditation and sleepiness during the meditation period, pre-meditation SMS scores and mean SSS ratings were included as covariates. Specifically, to isolate the change in mindful awareness during meditation practice, we regressed out pre-session SMS scores ($Ysmspre_t$), mean SSS ratings ($Ysss_t$), age ($Yage_t$) and sex ($Ysex_t$) from the post-session SMS scores ($Ysmspost_t$) at each time point $t$, thereby removing variance related to baseline mindful awareness, sleepiness and participant-specific characteristics from each session (eq. 1).

$$Yres_t = Ysmspost_t - (\beta_{1,t} Ysmspre_t + \beta_{2,t} Ysss_t + \beta_{3,t} Yage_t + \beta_{4,t} Ysex_t) \quad --(1)$$

$\beta_{1-4,t}$ denote parameters of the covariate predictors in eq. (1).

Using the residuals ($Yres_{1-6}$) from eq. (1), we performed repeated measures ANCOVAs for each participant across the six timepoints, with time since baseline ($T_{1-6}$) as the predictor variable (eq. 2).

$$\begin{bmatrix} Yres_1 \\ Yres_2 \\ Yres_3 \\ Yres_4 \\ Yres_5 \\ Yres_6 \end{bmatrix}_p = \beta_0 + \beta_{1,p} * \begin{bmatrix} T_1 \\ T_2 \\ T_3 \\ T_4 \\ T_5 \\ T_6 \end{bmatrix}_p \quad --(2)$$

$\beta_0$ denotes the parameter of the constant predictor, and $T_t$ denotes time elapsed between baseline (time point 1) and time point $t$.

This enabled us to examine whether mindful awareness during meditation increased proportionally to the time elapsed since the baseline meditation session, assuming a linear proportional relationship between time elapsed and mindful awareness. The resulting regression slope ($\beta_{1,p}$) from each participant $p$ was entered into a one-way Analysis of Variance (ANOVA) to examine group differences. This approach was followed independently for each SMS subscale, i.e., SMS-Mind and SMS-Body.

##### Change in breath counting performance

We evaluated changes in breath counting performance from baseline to the end of the weeklong post-NF self-guided meditation practice, using BCT task and probe accuracies. Difference scores of accuracies (follow-up minus baseline) of BCT task performance and probing were calculated. ANCOVA was used to detect significant group differences in task accuracy difference scores, with age and gender as covariates. Group differences in median probe accuracy difference scores were assessed using the non-parametric Wilcoxon rank sum test. In addition to the two participants excluded from behavioural analyses (as aforementioned), one participant's BCT data was not saved, resulting in N=37 participants (N=18 experimental, N=19 control) for this analysis.

False-discovery Rate (FDR) correction at a level of 5% was applied to control across all the five behavioural tests (**Table S2**). Effect sizes were estimated by transforming each resulting t-statistic to Cohen's *d.*

#### Offline fMRI data analysis

##### Change in NF target during NF-guided meditation: Activation

Activation in the NF target ROI (PCC) during meditation relative to rest was estimated offline through GLM in FSL FEAT (https://web.mit.edu/fsl_v5.0.10/fsl/doc/wiki/FEAT.html; FSL v6.0.6.4). The canonical double-gamma hemodynamic response function (HRF) was convolved with the time course of each fMRI condition (meditation, rest, and NF, as shown in **Figure 1B**) in the NF runs. The remaining cue/instruction condition was implicitly modelled by the constant term in the GLM. The HRF convolved time courses served as block design predictors in a first-level GLM to model the voxel-wise BOLD time series within the PCC in the smoothed preprocessed fMRI data from each NF run. The GLM also included nuisance regressors for MRI artefacts, head motion, and physiological noise, as described under 'Offline MRI preprocessing'. Parameter estimates (beta values) from the GLM fit were computed for the contrast of meditation vs. rest.

These parameter estimates were entered into second-level GLM in FSL FEAT to calculate the voxel-wise average of PCC beta values across the NF runs within each session for each participant. The mean framewise displacement (mFD), quantifying overall head motion in each run, was included as a covariate in the second-level GLM. Group differences in the mean residualised PCC beta values during each NF session were evaluated using third-level GLMs (ANCOVA) in MATLAB, with age, sex and average SSS ratings as covariates.

#### Change in NF target during NF-guided meditation: Functional Coupling

The neurocognitive model of focused attention meditation (*6*) suggests that deactivations in the DMN, associated with mental activity, are typically accompanied by activations in the SN and CEN, which support interoceptive awareness and cognitive control respectively. This mechanism likely reduces mental activity, enabling better focus on and awareness of the breath. Based on this model, we aimed to examine changes in context-dependent functional decoupling (i.e., negative correlation between regions during a state) between the NF target ROI (PCC - a core node of the DMN) and all voxels within the SN and CEN during NF-guided meditation vs. rest, using generalised psychophysiological interactions (gPPI) analysis (*39*) in FSL FEAT and Permutation Analysis of Linear Models (PALM) (*82*). The implementation of gPPI followed the GLM approach, with block design predictors in the first-level gPPI modelling the psychophysiological interactions between the seed region (PCC) and all voxels within CEN and SN. CEN and SN were outlined using the standard 7-network parcellation (*83*), applied to a grey matter mask created by averaging all participant-level masks and thresholding at 0.3. Similar to the GLM model described earlier, nuisance regressors capturing MRI artefacts, head motion, and physiological noise were included as covariates.

In the first-level GLM modelling for gPPI, the double-gamma HRF function was convolved with three block design time series: meditation minus rest, meditation plus rest, and NF condition, resulting in three condition predictors. The 'meditation minus rest' predictor captures the contrast of meditation > rest, isolating the differences between meditation and rest while excluding shared variance captured by the 'meditation plus rest' predictor. Although 'meditation plus rest' and 'meditation minus rest' span the same vector space as meditation and rest individually, the former approach directly isolates the desired contrast of meditation > rest. The predictors were zero-centred and convolved with the mean PCC BOLD time series (seed time course) to form the PPI predictors. Overall, the first-level model included the three condition predictors, their respective PPI predictors and the seed time course as the non-nuisance regressors, thereby spanning the entire experimental space (*39*). Beta maps for the PPI predictor of meditation > rest were generated from each NF run.

These beta maps were entered into second-level GLM to calculate the average voxel-wise beta values across NF runs within each NF session, with run-wise mFD values included as covariates. Subsequently, group differences in the residualised beta maps of each NF session were assessed using a third-level GLM in PALM, with age, sex and average SSS ratings as covariates. The resulting statistical maps indicating group differences in each session were thresholded at a cluster-forming threshold of $p<0.005$, and subjected to non-parametric permutation testing (10,000 permutations) with family-wise error (FWE) correction (FWE $p<0.05$) for multiple comparisons across clusters. Using this approach, we identified brain clusters within the SN and CEN that exhibited significant group differences in their extent of functional decoupling from the NF target during NF-guided meditation vs. rest.

##### Change in NF target from baseline to transfer meditation: Functional Coupling

We examined changes in functional coupling from pre-NF baseline meditation to post-NF transfer meditation between the NF target (PCC) and the significant NF gPPI cluster/(s). No NF was provided during the baseline and transfer tasks. Functional coupling was estimated here through Pearson's correlation (i.e., functional connectivity (FC)) between the significant gPPI cluster/(s) and PCC during the baseline meditation task and transfer meditation task. Difference scores were computed by subtracting baseline FC from transfer FC. Group differences in these FC difference scores were analysed using ANCOVA, with mean SSS, mFD, age, and sex as covariates.

#### Offline brain-behaviour association analysis

We analysed the relationship between significant NF target changes (PCC activation and/or PCC-seeded gPPI cluster) and significant behavioural outcomes (DASS-21, SMS slopes, and/or BCT accuracy) using Pearson's partial correlation, controlling for age and sex. Variance from mean SSS and mFD was regressed out from neural measures where applicable. For significant correlations, post-hoc group-wise correlations were performed to explore which group drove the observed sample-level correlations.

A 5% False Discovery Rate (FDR) correction was applied to control for multiple comparisons across the six PCC-based analyses not involving voxel-wise testing (i.e., the ANCOVAs and brain-behaviour correlations; **Table S2**). Where applicable, t-statistics were transformed to Cohen's *d* effect sizes.

#### Control analyses

##### Group assignment blinding

We assessed whether group blinding was effectively maintained during the NF training by evaluating differences in participants' post-NF self-ratings on the utility of NF for learning meditation, on in-scanner meditation performance, and on the correspondence between NF scores and meditation. We used non-parametric Wilcoxon rank sum tests to examine group differences in median ratings for each NF session on these three questions.

We also assessed the correlation between the sham and actual mean PCC signals associated with the control group in each NF session, using both online and offline denoised signals.

Additionally, we evaluated group differences in all the assessments and characteristics measured at baseline, including age, sex, meditation experience (hours), dispositional mindfulness (FFMQ and sub-scales), dispositional mind-wandering (MWQ), dispositional anxiety (STAI-T), 1-month sleep quality (PSQI), 1-week emotional distress (DASS-21), and breath counting (BCT task accuracy). Two-sample t-tests were used for all the group comparisons, except for sex and ethno-geographic background, where Chi-square tests were used to compare proportions between groups.

##### Association between online and offline PCC signals

We assessed the validity of the PCC signal calculated online during NF sessions by comparing it to the PCC signal estimated offline after thorough denoising and preprocessing. This was done by examining the correlation between each session's mean offline denoised PCC beta values and mean online denoised PCC PSC.

##### Group differences in online physiological proxy

We explored whether the online activation in the control ROI, a proxy for online physiological fluctuations, differed significantly between groups during NF-guided meditation vs. rest. ANCOVAs were conducted to assess session-wise group differences in the confound ROI's mean PSC during meditation relative to resting baseline, adjusting for age and sex as covariates.

### Supplementary Section 2

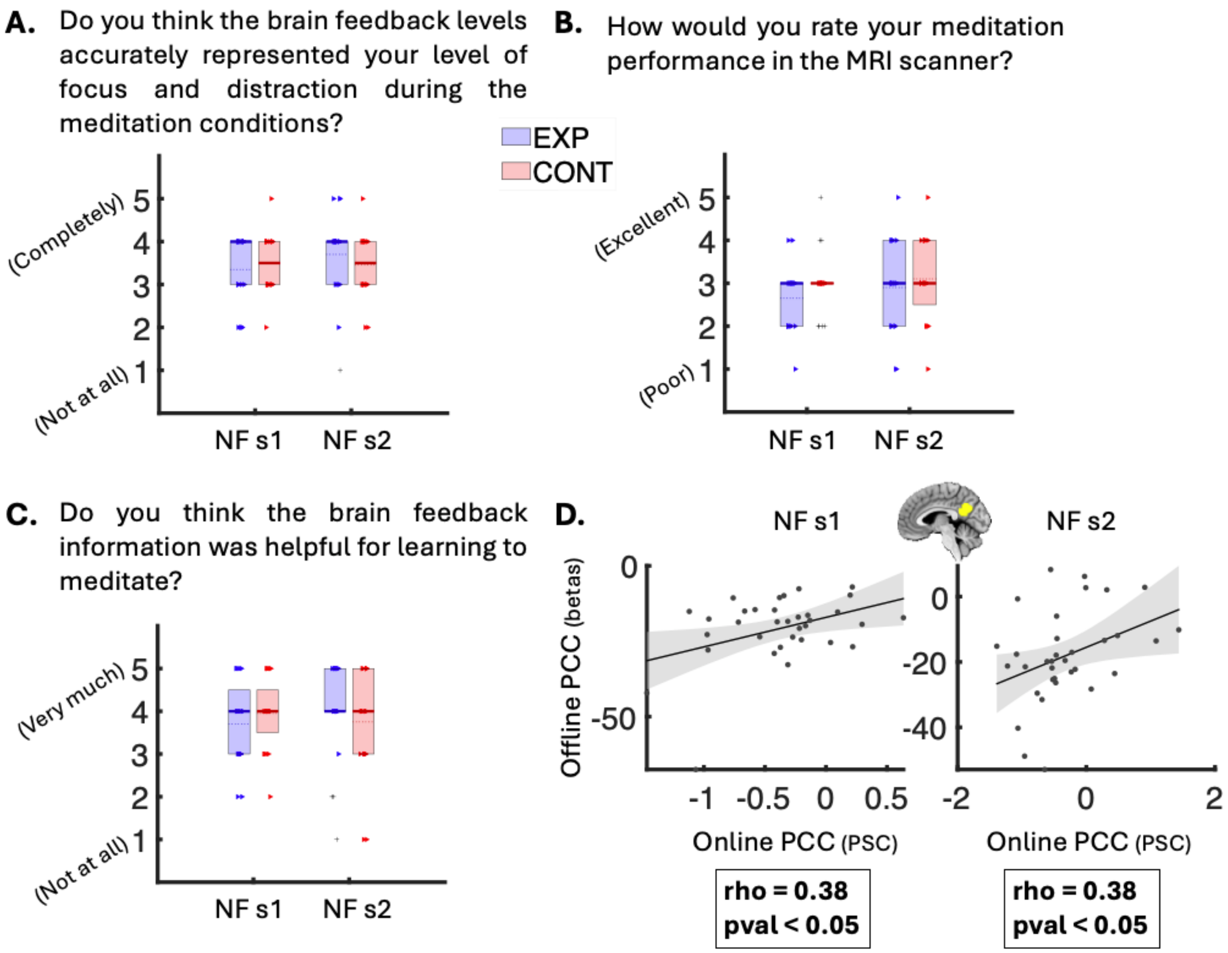

***Figure S3:*** *Assessment of blinding effectiveness and the relationship between online and offline PCC signals.* ***A) to C)*** *Box plots depicting participant ratings (1 to 5 scale) for each question indicated above the plots, collected at the end of each NF session. Blue represents the experimental group, and red represents the control group. No significant between-group differences in median ratings were observed for any of the questions (p > 0.05). Triangles indicate individual data points, with solid lines marking medians, dashed lines marking means, and pluses indicating outliers.* ***D)*** *Scatter plots displaying significant positive correlations between PCC activation betas denoised post-hoc (y-axis) and PCC activation PSC (x-axis) denoised during NF, for the contrast meditation vs. rest across the two NF sessions (s1: p = 0.028, rho = 0.38; s2: p = 0.028, rho = 0.38; N=34). Dots represent individual data points, with the shaded area showing the 95% confidence interval around the linear fit (solid black line). NF - neurofeedback; s1 - session*

*1; s2 - session 2; EXP - experimental group; CONT - control group; PCC - posterior cingulate cortex; PSC - percent signal change*

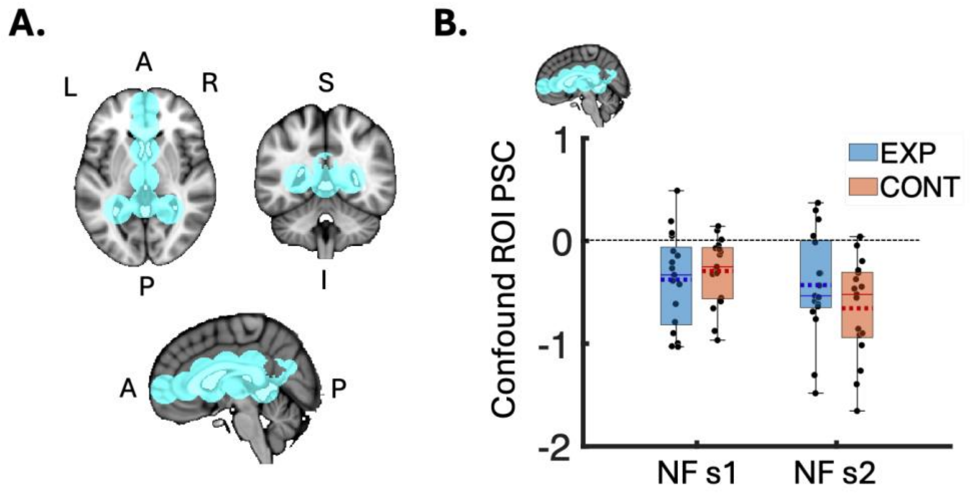

***Figure S4:*** *Control region of interest (ROI) used during NF as proxy for online denoising, and its percent signal change (PSC).* ***A)*** *Brain volume slices depicting the anatomical extent of the control ROI (in cyan) used as proxy for online physiological denoising during NF training.* ***B)*** *Box plot of mean PSC estimated online from the confound ROI during each NF session. There were no significant differences between groups in either session ($p > 0.05$; N(exp) = 17, N(cont) = 17). The box plot shows individual data points as black dots, outliers as triangles, data range as whiskers, means as dotted lines, and medians as solid lines. NF - neurofeedback; s1 - session 1; s2 - session 2; ROI - region of interest; PSC - percent signal change; EXP - experimental group; CONT - control group; P - posterior; L - left; R - right; I - inferior; S – Superior*

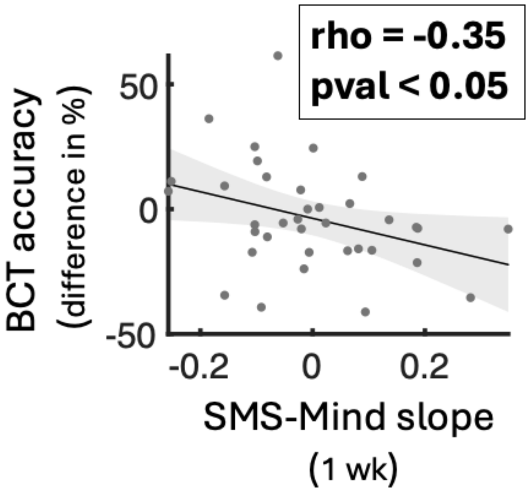

***Figure S5:*** *Scatter plot of the post-hoc correlation analysis illustrating a significant negative correlation between increase in BCT accuracy from baseline to follow-up (y-axis; difference in BCT accuracy %) and decrease in slopes of mindful awareness of mental activity (x-axis; SMS-Mind slopes) (p = 0.044; rho = -0.35; N = 36). Individual data points are represented by dots, with the shaded area indicating the 95% confidence interval around the linear fit (solid black line). BCT - breath counting task; SMS - state mindfulness scale; wk - week*

**Table S2**: Summary of findings with key statistics

| | **Analysis summary** | **Cohen's *d* effect size OR correlation coefficient *rho* (dfe)** | **FDR-adjusted p value (Uncorrected p value)** |
|---|---|---|---|
| **Behaviour** | 5% FDR correction across all five behavioural tests below | | |

| | | | |
|---|---|---|---|
| Change in emotional distress | ANCOVA group difference in total *DASS-21* difference scores | 0.40 (34) | **0.041** (0.026) |
| Rate of change in mindful awareness of *mental activity* during real-world meditation | ANCOVA group difference in 1-week *SMS*-Mind slopes | 0.38 (36) | **0.041** (0.027) |
| Rate of change in mindful awareness of *bodily sensations* during real-world meditation | ANCOVA group difference in 1-week *SMS*-Body slopes | 0.00 (36) | 0.988 (0.988) |
| Change in breath counting skill | ANCOVA group difference in *BCT* task performance accuracy | 0.39 (32) | **0.041** (0.033) |
| | Wilcoxon rank sum test group difference in *BCT* probe accuracy | 0.56 (34) | **0.005** (0.001) |
| **Brain** | | | |
| **Voxel-based** | 5% FWE correction with 10,000 permutations | | |
| Change in PCC-seeded decoupling during NF meditation vs. rest | ANCOVA group difference from voxel-based gPPI analysis (NF session 1) | - | n.s. |
| | ANCOVA group difference from voxel-based gPPI analysis (NF session 2) | DLPFC cluster; 0.59 (29) | *FWE*-adjusted p = **0.032**; cluster size = 26 voxels |
| **ROI-based** | 5% FDR correction across all six ROI-based tests below | | |
| Change in PCC activation during NF meditation vs. rest | ANCOVA group difference (NF session 1) | 0.04 (29) | 0.876 (0.814) |
| | ANCOVA group difference (NF session 2) | 0.03 (29) | 0.876 (0.876) |
| Change in PCC-DLPFC FC from baseline to transfer meditation without NF (session 2 due to gPPI significance in session 2) | ANCOVA group difference in FC difference values | 0.14 (28) | 0.715 (0.477) |
| **Correlations** | | | |
| Correlation between PCC-DLPFC decoupling (NF meditation session 2) and change in emotional distress | Pearson's partial correlation, controlling for age and gender | 0.53 (28) | **0.018** (0.003) |
| Correlation between PCC-DLPFC decoupling (NF meditation session 2) and slope of change in mindful awareness of mental activity | Pearson's partial correlation, controlling for age and gender | -0.14 (28) | 0.715 (0.462) |
| Correlation between PCC-DLPFC decoupling (NF meditation session 2) and change in breath counting accuracy | Pearson's partial correlation, controlling for age and gender | -0.23 (27) | 0.660 (0.220) |

FDR - Benjamini-Hochberg false discovery rate; FWE - family-wise error; n.s. - not significant; dfe - degrees of freedom; ANCOVA - analysis of covariance; DASS-21 - depression, stress, anxiety scale; SMS - state mindfulness scale; BCT - breath counting task; NF - neurofeedback; PCC - posterior cingulate cortex; DLPFC - dorsolateral prefrontal cortex; ROI - region of interest; gPPI - generalised psychophysiological interactions; FC - functional connectivity

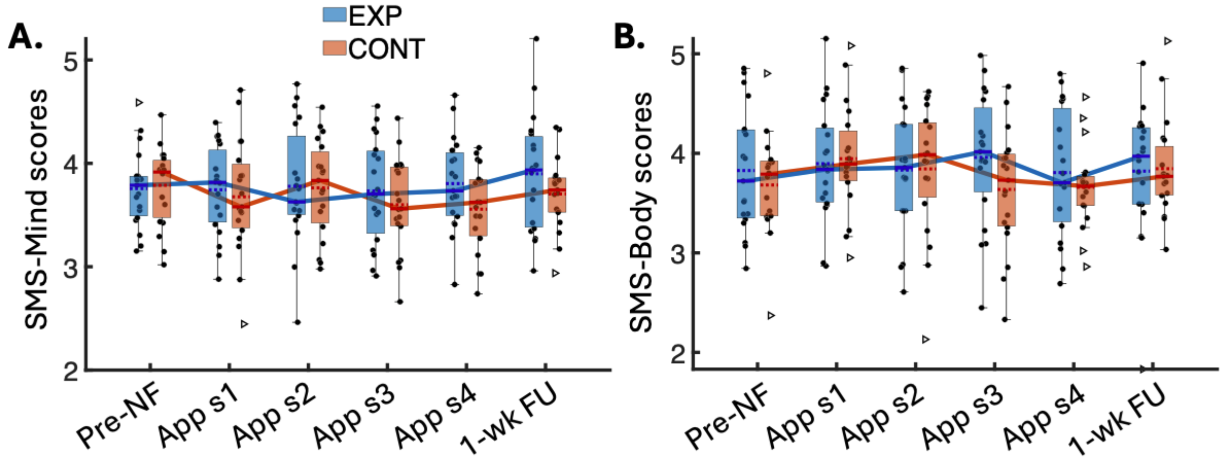

***Figure S6**: Visualisation of changes in mindful awareness (SMS subscale scores) during 5-minute self-guided meditation sessions over one week post-NF. **A)** Box plot showing changes in mindful awareness of mental activity (SMS-Mind) over time in each group, adjusted for pre-meditation SMS-Mind, sleepiness, age, and sex. **B)** Similar plot for mindful awareness of bodily sensations (SMS-Body). Dots represent individual scores, triangles denote outliers, whiskers indicate data range, and the solid and dashed lines within each box represent the median and mean, respectively. Medians across time points are connected to illustrate trends, with blue for the experimental group and red for the control group. NF - neurofeedback; sx - session x; EXP - experimental group; CONT - control group; FU - follow up; SMS - state mindfulness scale; wk - week*
